# Filtering for highly variable genes and high quality spots improves phylogenetic analysis of cancer spatial transcriptomics Visium data

**DOI:** 10.1101/2024.07.11.603166

**Authors:** Alexandra “Sasha” Gavryushkina, Holly R Pinkney, Sarah D Diermeier, Alex Gavryushkin

## Abstract

Phylogenetic relationship of cells within tumours can help us to understand how cancer develops in space and time, identify driver mutations and other evolutionary events that enable cancer growth and spread. Numerous studies have reconstructed phylo-genies from single-cell DNA-seq data. Here we are looking into the problem of phylogenetic analysis of spatially resolved near single-cell RNA-seq data, which is a cost-efficient alternative (or complementary) data source that integrates multiple sources of evolutionary information including point mutations, copy-number changes, and epimutations. Recent attempts to use such data, although promising, raised many methodological challenges. Here, we explored data-preprocessing and modelling approaches for evolutionary analyses of Visium spatial transcriptomics data. We conclude that using only highly variable genes and accounting for heterogeneous RNA capture across tissue-covered spots improves the reconstructed topological relationships and influences estimated branch lengths.

## 1 Introduction

Reconstructing single cell phylogenies within tumours can shed light on cancer development (Nowell, 1976) and help identify driver mutations and other evolutionary events that enable cancer growth and spread (Koch, 2022). Single cell phylogenies have been reconstructed from different types of data: single cell DNA data (Navin et al., 2011), induced mutations data (Baron and van Oudenaarden, 2019), barcode data (McKenna and Gagnon, 2019), and recently from single cell RNA-sequencing data (Moravec et al., 2023; Liu et al., 2023). In addition to the evolutionary signal conveyed through mutations, single-cell RNA-sequencing data captures expression levels influenced by other evolutionary changes, such as copy number variation or changes in epigenetic and transcriptional regulation mechanisms. Encompassing these different aspects of cell’s evolution, such data could prove to be more cost-effective for phylogenetic reconstruction.

To date, there have been very few attempts to reconstruct single cell phylogenies from single cell RNA data. Moravec et al. (2023) attempted to reconstruct single cell phylogenies of cancer using either single nucleotide variant (SNV) or expression data obtained using droplet based single cell RNA-sequencing techniques. Liu et al. (2023) used an imputation technique to reconstruct cell phylogenies of cancer from sparse SNV data. Other studies used expression data to reconstruct phylogenetic trees of cell types within and between species (Liang et al., 2015; Mah and Dunn, 2024).

Although promising, phylogenetic reconstruction from expression data presents several challenges, that can be categorised in three groups. First of all, there are challenges connected to the technology limitations, e.g., low detection efficiency (Hicks et al., 2018), uneven RNA capture across cells (Hafemeister and Satija, 2019), multicellular resolution in the case of Visium Spatial Transcriptomics (ST) data (Moses and Pachter, 2022), sequencing errors, ambiguous mapping, among others.

Second, biological function of cells and their environment influence expression of many genes and using such genes in a phylogenetic analysis might be misleading. Therefore, selecting genes where expression reflects evolutionary history rather than biological function or environmental impact is crucial for estimating reliable single cell phylogenies.

Last, biological, statistical and computational problems connected to modelling of expression evolution need to be addressed. Evolutionary change in the expression levels can be modelled as a Brownian motion (Felsenstein, 1973) or Ornstein–Uhlenbeck processes (Hansen, 1997; Dimayacyac et al., 2023), however when a large number of genes is used in a phylogenetic analysis, such models become computationally expensive and prohibitive. One way to address this problem is discretising expression counts (Moravec et al., 2023) and using ex-isting models of morphological (Lewis, 2001) or molecular (nucleotide or amino acid) character evolution Tavaŕe and Miura (1986). Additionally to statistically assessing different models performance, these different approaches can be assessed based on the biological interpretability of the obtained phylogenies.

Spatially resolved, near single-cell RNA sequencing techniques have become available (Asp et al., 2020; Moses and Pachter, 2022) and provide new opportunities for studying gene expression in the context of the 2D organisation of a tissue, for example to investigate cancer evolution. Single cell phylogenies reconstructed from such data can shed light on spatial development of tumours using phylogeography (Lemey et al., 2009; Bouckaert, 2016) and phylodynamics methods (Lewinsohn et al., 2023). Phylogeography methodology is particularly promising here as it links genomic variability and evolution to population organisation and dynamics in space. In the context of cancer, this would allow studying such fundamental biomedical question as genomic mechanisms of metastatic disease and treatment resistance development by, for example, performing ancestral state reconstruction of tumour cells in space and time.

However, the reliability of different approaches to reconstructing single cell phylogenies from such data has not been investigated. Here we explored approaches for phylogenetic analysis of Visium ST expression data. We conclude that gene expression variability across cells and uneven RNA capture across tissue covered spots has a noticeable impact on the reconstructed phylogenetic relationships of cells.

## 2 Results

The ST data analysed in this study have multicellular resolution with each capture spot covering several cells. In what follows, we assume that cells under the same capture spot are genetically similar and each spot covers approximately the same number of cells. Having these assumptions, we interpret phylogenetic trees reconstructed from multicellular spots as single cell phylogenetic trees.

We analysed Visium ST data from two patients with colorectal cancer from eight sections across five tissues (Supp. Figure 8). We reconstructed phylogenetic trees relating spots from primary tumour, normal colon and matched metastatic tissue with enough expression (referenced as 50% sample) using a maximum likelihood approach (iqtree2 Minh et al. (2020)). We also performed Bayesian phylogenetic analyses (BEAST2 Bouckaert et al. (2019)) on subsampled data, using one subsample from patient 1 (referenced as similar quality sample) and four subsamples from patient 2 (referenced as test, highest quality, similar quality, and manually selected samples). Subsamples are described in Methods 4.3. For each of the subsamples, we performed four analyses with different pre-processing, modelling and gene-filtering approaches. We compared normalised vs unnormalised expression counts; discretised expression counts assuming an ordinal model vs raw or continuous expression counts assuming a Brownian motion model, and all expressed genes vs highly variable genes (HVGs) only (Methods 4.2-4.5).

The maximum likelihood trees are shown in figure 1. The patient 1 tree displays a normal clade consisting of spots from two normal colon sections and a malignant clade that consists of spots from two tumour sections with a small subclade of twelve normal spots. The patient 2 tree has two major malignant clades and one major metastatic clade, however there is a certain degree of intermixing of primary and metastatic samples within these clades. Additionally, there is a separate metastatic clade that diverges within one of the major primary malignant clades and a primary malignant clade that diverges within a major metastatic clade. Both trees show very long external branches compared to the internal branches. In both trees and other analyses, we observed good mixing of samples from serial sections (Supp. Figure 32).

**Figure 1:**
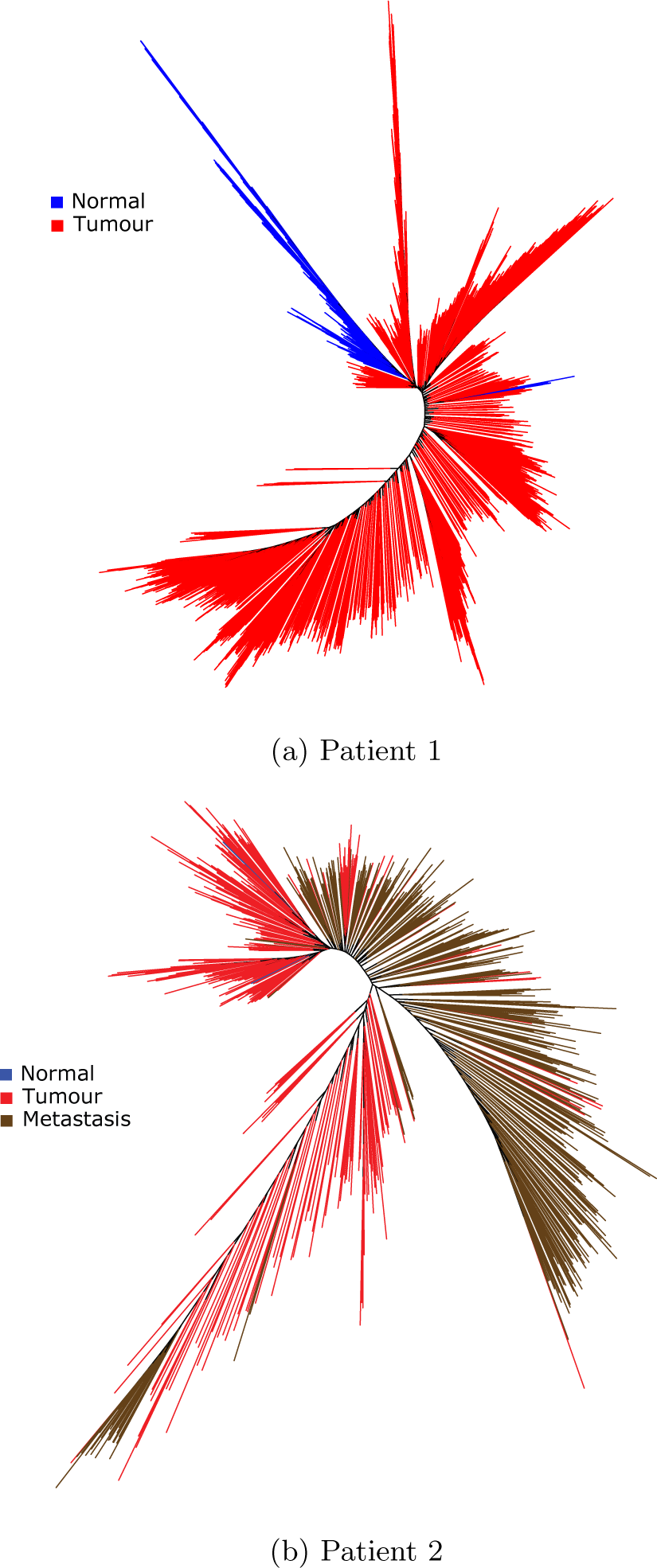
Maximum likelihood trees of 50% density samples of patient 1 (a) and patient 2 (b). (a) In the tree for patient 1, there is a major malignant and a major normal clade as expected. However, there is also another normal clade nested within the malignant clade, which is a biologically implausible scenario. (b) There are only two normal spots in 50% density sample of patient 2. Overall there are no clades that consist primarily of the spots of the same type however there are three noticeable clades, one comprised of mostly metastatic spots and two comprised of mostly primary malignant spots. There are also two noticeable subclades within two of the major clades: a metastatic subclade that nests within one of the major primary clades and a primary subclade that nests within the major metastatic clade, which are again biologically implausible scenarios. The unexpected phylogenetic relationships in both patients can be attributed to the limitations of technology, data pre-processing, variable nature of expression data, inadequate modelling approaches, or inaccuracies in the representation of the probable phylogenetic relationships by a single tree.

To assess the performance of the different approaches to the phylogenetic analysis of Visium ST expression data we looked at how well reconstructed phylogenies describe expected evolutionary history of spots from different sections and tissue types. Spots on all sections from both patients were annotated by a pathologist (Pinkney et al., 2024) and could be broken into five major tissue types: malignant epithelium, malignant stroma, benign epithelium, benign stroma and benign lymphocytic aggregation (Supp. Figure 10). The tumour sections have both malignant and benign adjacent normal tissue types in both patients. The metastatic section from patient 2 has only malignant gland tissue (desmoplastic stroma with low cellularity was not analysed).

We sampled spots for Bayesian analyses accounting for the tissue types. For the test sample, we selected spots from two malignant gland regions and one malignant stroma region of patient 2 tumour 2 section (Supp. Figure 12). We expected that the three regions will form clades in the reconstructed phylogenies. However, no clear separation of the regions was noticed in any of the summary trees of the four performed analyses (Supp. Figures 13, 14, 15, 16).

To assess whether we can reconstruct the evolution of the normal tissue into cancerous, we performed several Bayesian analyses of benign epithelium and malignant epithelium from all sections including adjacent normal benign epithelium from tumour sections. Accounting for the sections, we have three section/tissue types for patient 1: benign epithelium from normal sections, benign adjacent normal epithelium from tumour sections, malignant epithelium from tumour sections. For patient 2 we have the same three categories, with an additional category for metastatic malignant epithelium from the matched metastasis section. Our expectation was that spots of similar tissue/section types would form clades and display certain evolutionary relationships. In particular, we expected that the most recent common ancestor (the root of the phylogeny) was a normal cell and in accordance with the parsimony principle, the number of times a normal lineage evolved into a malignant lineage or a primary malignant lineage evolved into a metastatic lineage was small. We measured the plausibility of the reconstructed phylogenetic relationship by the number of switches from normal to malignant lineage (number of malignant clades), or from primary malignant or normal to metastatic lineage (number of metastatic clades) observed in such phylogenies. Note that we do not see intermediate stages of lineages in the reconstructed phylogenies, which implies that a normal to metastatic switch could be through an unobserved intermediate primary malignant stage of the lineage Leung et al. (2017); Chen et al. (2022a) or could represent an early metastatic seeding Hu et al. (2019).

Overall the best result in terms of the average number of malignant and metastatic clades in the posterior distributions of trees was achieved for the analysis of discretised counts of HVGs (discretised HVGs) (Figure 6 a, b). Similar performance was achieved when normalised and discretised counts of HVG were analysed. Using discretised counts of all genes or continuous HVGs resulted in similar relatively poorer performance.

For patient 1, reconstructed phylogenies satisfied our expectations in all analyses, Figure 2 and Supp. Figures 18, 19, 20. Patient 2 data were particularly challenging for the phylogenetic analysis and reconstructed trees met our expectations to a lesser degree. The discretised HVGs analysis of the high quality sample displays expected evolutionary relationship except for a single misplaced benign tumour branch in the maximum clade credibility (MCC) tree (Figure 3) and displays the smallest number of malignant and metastatic clades (Figure 6a and b). However, such near perfect evolutionary relationships could be attributed to the prior use of the annotation for selecting spots. The highest quality spots were chosen separately within each tissue/section type resulting in high differences in total counts for spots of different types. The analyses of the similar quality subsample, where cells with similar total counts were chosen, produced a tree with high uncertainty (Figure 4). Out of the clades with relatively high supports (*≥*0.22), we could distinguish a malignant clade with a metastatic clade diverging within it and a clear normal clade. There is however a clade that contains a mix of benign and malignant spots from tumour sections and the normal clade unexpectedly diverges within this mixed clade. One of the normal spots is recovered as sister to the rest of the spots with probability 0.96. The MCC tree for the HVG discretised analysis of manually selected spots (in which the quality of spots can be considered random, Figure 5) displays some of the expected evolutionary relationship with a clear benign and two malignant clades, with one of the malignant clades diverging within the benign clade. Unexpectedly, two metastatic and one primary malignant spots are recovered as sister to the normal clade.

**Figure 2:**
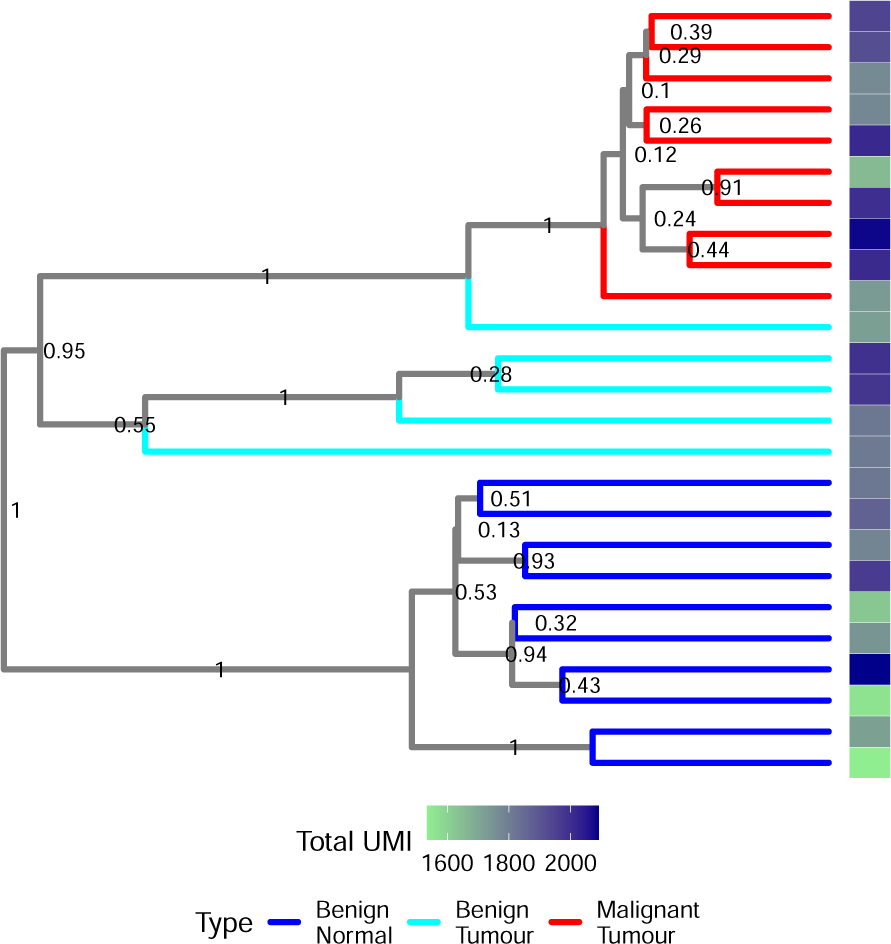
MCC tree of a BEAST2 analysis of similar quality sample from patient 1 with raw discretised counts for HVGs. Colours reflect the tissue type and numbers are posterior support values for clades. Bar on the right shows total UMI counts of spots. The tree shows expected evolutionary history of benign epithelium spots from normal and tumour section and malignant epithelium spots. The support values are relatively high, in particular, at the tissue/section type divergences. There is no obvious correlation between phylogenetic distance and total UMI counts.

**Figure 3:**
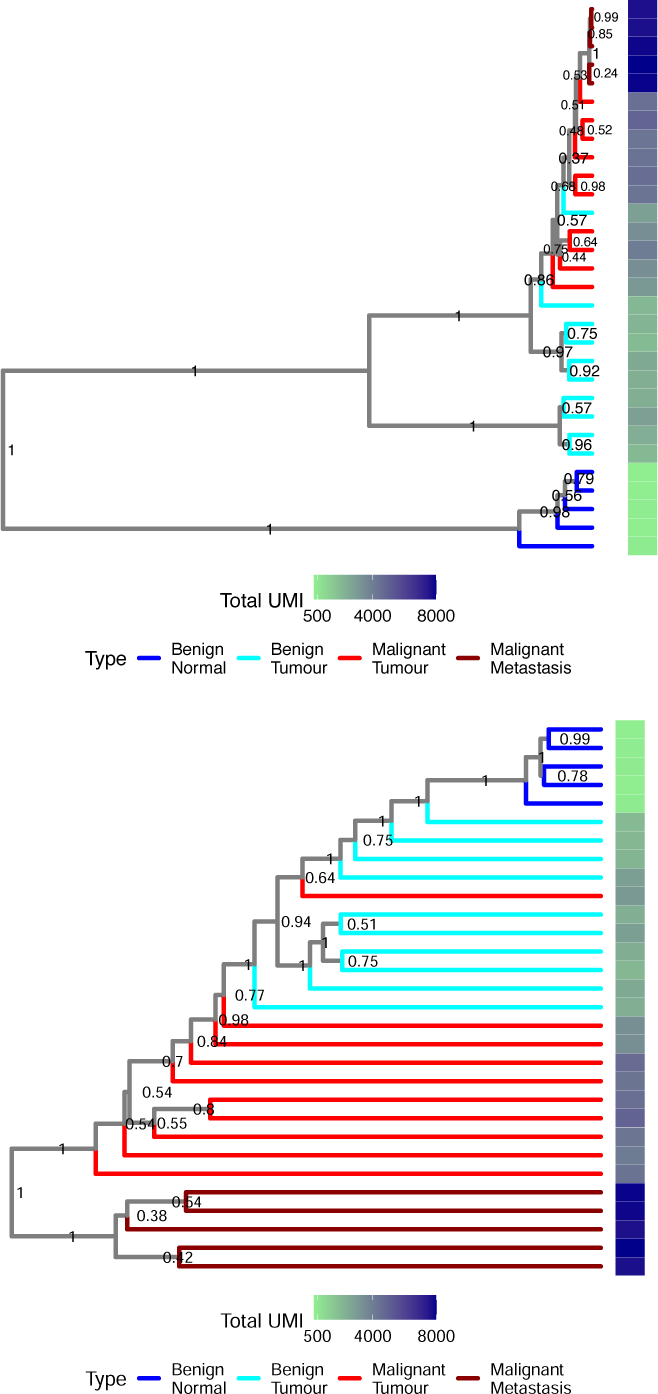
MCC trees of two BEAST2 analyses of highest quality sample from patient 2 with discretised counts for HVGs (top) and all genes (bottom). Colours reflect the tissue type and numbers are posterior support values for clades. Bars on the right show total UMI counts at each spot. The discretised HVGs tree (top) shows plausible evolutionary scenario where benign Tumour clade diverges from benign Normal, a malignant clade diverges from benign Tumour, and a metastatic clade diverges within a malignant primary clade. Although the tree reconstructed using all genes keeps the tissue/section type separation, it either displays an unrealistic divergence of the benign epithelium clade within the malignant clade or can be interpreted as having an unrealistically large number of malignant clades. When HVGs are used, the evolutionary history reflects our expectations with malignant epithelium diverging within adjacent normal epithelium clade and metastatic malignant epithelium diverging within primary malignant epithelium clade. The support values are overall high with many clades having 100% supports in both trees. There is a noticeable correlation between phylogenetic distance and total UMI counts. External branch lengths are considerably longer relative to the tree length in the tree reconstructed from all genes.

**Figure 4:**
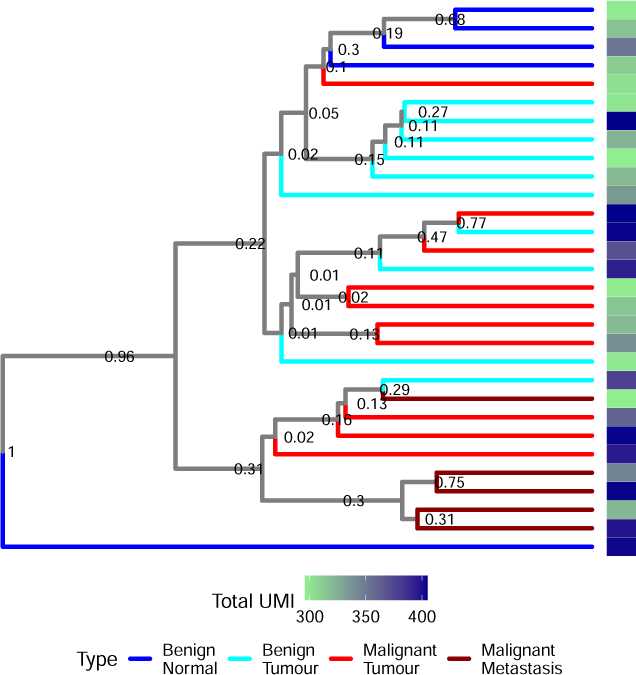
MCC tree for the BEAST2 analysis of the similar quality sample from patient 2 with discretised counts of HVGs. Colours reflect the tissue/section type and numbers are posterior support values for clades. Bars on the right show total UMI counts at each spot. The tree displays a clear malignant clade consisting of three malignant spots from tumour sections and a metastatic clade diverging within this clade and a clear normal clade. The remaining malignant spots from the tumour sections form a clade with the benign spots from the tumour sections that also contains the benign normal clade. The support values are overall low. There is no obvious correlation between phylogenetic distance and total UMI counts.

**Figure 5:**
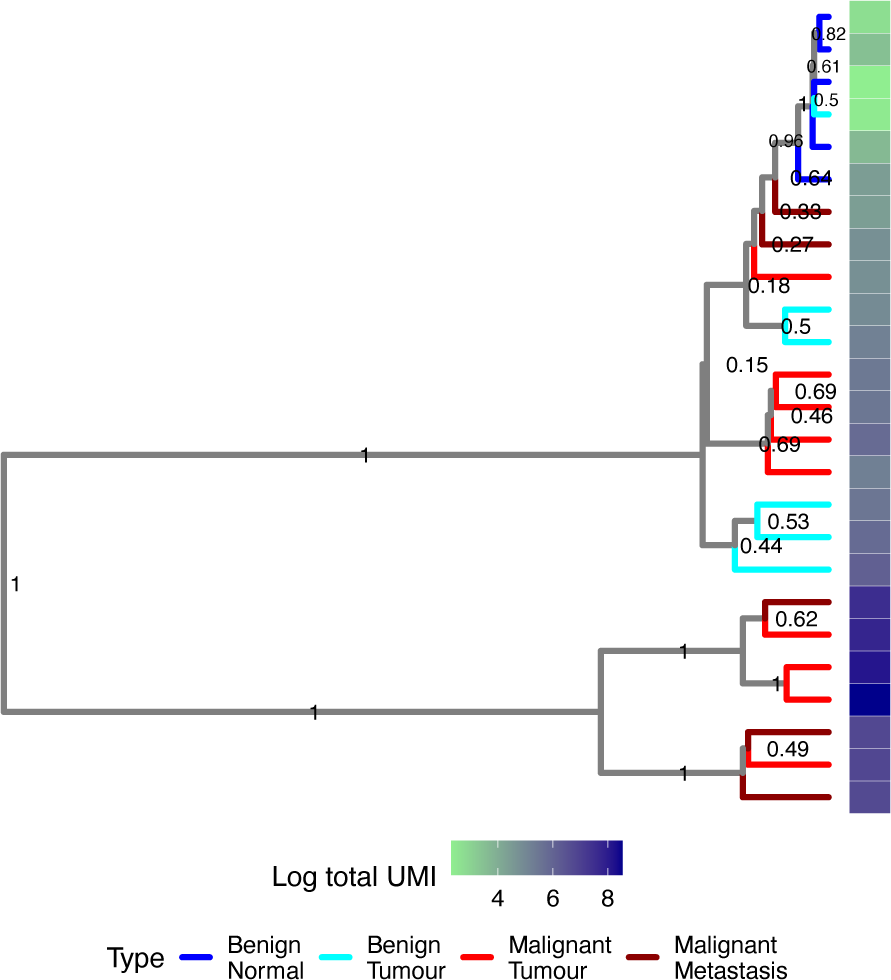
MCC tree for the BEAST2 analysis of the manually selected sample from patient 2 with discretised counts of HVGs. Colours reflect the tissue/section type and numbers are posterior support values for clades. Bars on the right show total UMI counts at each spot. The tree displays a clear benign normal clade, and two malignant clades. Metastatic spots do not form a clade with two metastatic spots together with one primary malignant placed as sister to the benign normal clade. The support values are overall high except for several deeper divergences within the clade consisting of all benign spots, one of the malignant clades, one primary malignant and two metastatic spots. There is a noticeable correlation between phylogenetic distance and total UMI counts.

Because archival tissue samples were used, there is a high level of variation in the total expression counts among spots within and between sections, (determined by evaluating unique molecular identifier (UMI) counts). Total counts variation remains high after the pre-processing SpaceRanger aggregation step where spot counts are downsampled to achieve the same average total UMI counts per spot on all sections (Supp. Figure 9). Overall normal sections have lower average total counts and standard deviations compared to tumour or metastatic sections but higher coefficients of variation (Table 2).

To account for such variation we either used downsampling to achieve the same average UMI counts per spot in the samples; normalised each spot expression counts by the spot total UMI count; or selected spots with highest (within tissue/section type) or similar total counts. Downsampling did not improve the expected phylogenetic relationship in several analyses (data not shown) potentially due to extreme data loss and was not assessed further. Normalisation did not change the average number of malignant or metastatic clades in posterior distribution of trees with maximum difference between analyses with normalized and unnormalised HVGs of 0.9. Overall, using normalised counts as compared to unnormalised counts produced trees (unnormalised analyses: Figures 2 3, 4, 5, and Supp. Figures 14; normalised analyses: Supp. Figures 15, 19, 23 26 30) with similar or less plausible phylogenetic relationships. Taking into account the quality of spots reflected by their total UMI counts had an effect on the number of malignant and metastatic clades as we described earlier (Figure 6). In test, manually selected, and high quality samples there is a positive correlation between the difference in the total UMI counts and phylogenetic distance of pairs of spots (Figure 7). However, when spots with similar total counts are selected very weak or no correlation is observed.

**Figure 6:**
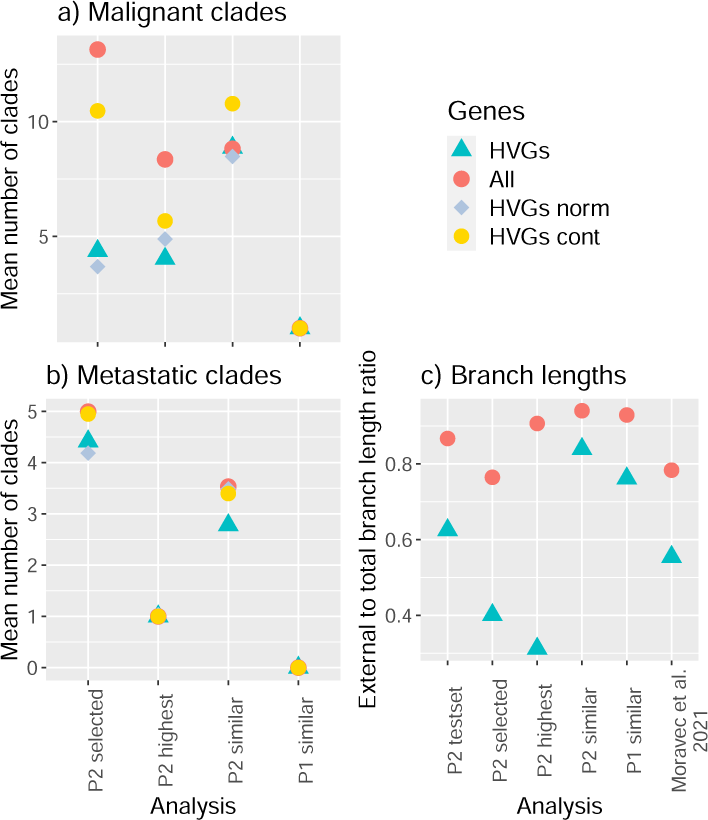
Mean number of (a) malignant (primary and metastasis) and (b) metastatic clades in posterior distributions of trees from three analyses of sam-ples from patient 2 (P2) and a similar quality sample from patient 1 (P1). (c) The ratio of the external to total branch lengths in MCC trees from Bayesian analyses of four samples from patient 2 (P2), similar quality sample from patient 1 (P1) and 50% density sample from Moravec et al. (2021).

**Figure 7:**
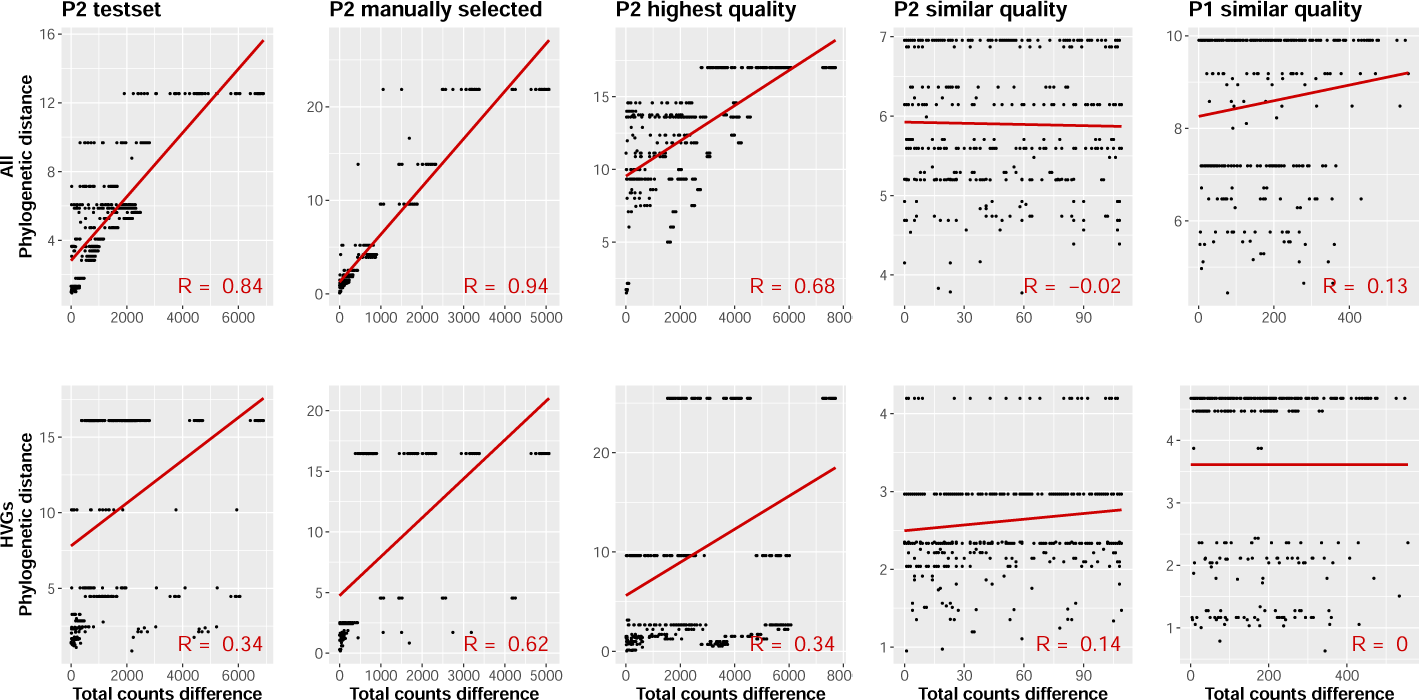
Phylogenetic distance against total counts difference for pairs of spots in each sample for analyses with discretised counts of all (top row) and HVGs (bottom row). Linear model regression lines and correlation coefficients (R) are shown in red. P1 = patient 1, P2 = patient 2.

Finally, we assessed how filtering for genes with higher expression variation among spots affects phylogenetic reconstruction. The MCC trees reconstructed using discretised counts of HVGs showed more plausible cancer development history compared to the analyses where all expressed genes in the sample were used, with smaller or equal numbers of malignant and metastatic clades in all four analyses (Figure 6 a, b). The most profound effect of using HVGs is found in the manually selected sample. Using HVGs also decreased the relative external branch lengths in all analyses including reanalysis of data from Moravec et al. (2023) (Figure 6 c).

## 3 Discussion

Here we explored the possibility of reconstructing cell phylogenies from expression data obtained by Visium ST technology. We attempted to address the problems of non-even RNA capture across spots, gene filtering and compare different approaches to modelling expression evolution. Overall we see that the discretisation approach proposed by Moravec et al. (2023) works well for Visium ST. We have detected two aspects that influence phylogenetic inference: the quality of sampled spots (as can be approximated by the total spot’s UMI counts) and gene expression variance.

Using HVGs is a common practice in expression data analysis. Here we hypothesised that genes that evolve neutrally and are potentially more likely to contain phylogenetic signal, would have higher variance in their expression values across spots. Phylogenetic analysis where only HVGs were used showed similar or more plausible evolutionary relationships in all considered cases. Using HVGs also resulted in the reduction of the unusually long external branches observed in the analyses where all or most expressed genes were used, including the original analyses of expression data in Moravec et al. (2023). Mah and Dunn (2024) noticed a similar effect when using only the first 20 principal components, which contain the most variation, in a phylogenetic analysis of between species expression data.

The observed variation of the spots’ total UMI counts in the analysed data could reflect both: the real differences in the expression of cells and/or the effects of the imperfect technology, e.g., differences in the detection efficiency across spots (spot quality) or unequal numbers of cells under spots. Normalising expression counts by the spots’ total UMI counts did not improve or only slightly improved (in one out of four analyses) the expected phylogenetic relationship. The downsides of such normalisation have been noticed in other analyses of single-cell RNA sequencing data (Church et al., 2022). Whereas, analysis where highest quality spots were selected showed the best results compared to the analysis where random quality (manually selected based on the location) or similar quality selected spots were analysed. In the analysis of manually selected spots, we observed two metastatic spots with lower total counts compared to other malignant spots (metastatic or primary), that are recovered as sister to the benign normal clade rather than any of the malignant clades. Such an unrealistic evolutionary position of the metastatic spots might be explained by the poor quality of these spots. These observations highlight the importance of selecting spots of high enough quality for a phylogenetic analysis.

The near perfect results of the highest quality sample analysis however can not be taken as the gold standard for the performance of the proposed approach, as it uses the spot annotation in a circular way, first for selecting high quality spots within the same tissue/section type and then for assessing the plausibility of the phylogenetic relationship. In the analysis of similar quality samples the expected phylogenetic relationships are recoverable to a lesser degree in patient 2, but are perfectly recoverable for patient 1. The data for patient 1 appears to be less challenging for phylogenetic reconstruction, as can also be seen in the maximum likelihood analyses where there is a clear separation of spots from normal and tumour sections in patient 1, whereas in patient 2 metastatic and primary tumour sections appeared less separable. Therefore, we focused on assessing the performance in the more challenging dataset. The similar quality analysis for patient 2 can be seen as an example of the worst case scenario when analysing a sample of similar and relatively high quality spots.

In the 50% density, highest, and similar quality analyses, malignant and benign clades contain spots from both serial sections and we do not observe clades comprised solely from samples of one of the serial section. This confirms that across section normalisation achieved by downsampling at the aggregation stage have eliminated the section effect. The manually selected sample analysis shows however that within section heterogeneity remains a problem and, as we have showed, can not be solved by simply normalising by total counts. Other more sophisticated normalisation approaches (Hou et al., 2020; Church et al., 2022) might improve phylogenetic reconstruction.

We also noticed correlation between the estimated phylogenetic distance and the difference in the total counts of pairs of spots even for analyses where only HVGs were considered. Since we observe a variation in spot’s total counts between sections and tissue types (Sup. Figure 9 and Figure 11), a certain degree of correlation is expected if we assume that spots of the same tissue/section types are phylogenetically close. However such relatively high correlation may imply that the phylogenetic reconstruction is driven primarily by the quality of the spots. The correlation subsides when similar total counts are considered and expected phylogenetic relationships remain recoverable in patient 1 however less recoverable in patient 2. The later could be attributed to the poorer quality of the data for patient 2, which is also reflected in the low support values for the majority of the clades in this tree. Low correlation for the similar quality sample in patient 1 suggests that there is phylogenetic signal in the expression data distinct from just the differences in spot qualities on different sections.

Here we assessed the performance of different approaches by how well the reconstructed phylogenies satisfied our expectation of cancer evolution. We considered a smaller number of normal to cancer evolutionary events supported by the reconstructed phylogenetic tree as an indicator of a potentially better approach. Some analyses produced unrealistically large number of malignant clades, primary malignant clades diverging within metastatic clades and otherwise misplaced branches. Such unlikely evolutionary relationships could be attributed to the poor phylogenetic signal in the expression data, limited data quality, and limitations of the approaches that we used.

First of all, although we attempted to address one of the technology limitations, namely, the uneven detection efficiency and poor spot quality through normalisation or selection of high quality spots, many other potential issues were not addressed. An important problem that we have not addressed in this study is the multicellular resolution of the Visium ST data. We made an assumption that the collection of RNA molecules captured at each spot approximates the expression of a single cell. Under this assumption, analysing spots that cover evolutionarily distant cells might produce unexpected results. The number of cells covered by a spot can also vary in a tissue-type dependent manner, which might further jeopardise phylogenetic reconstruction. Cell deconvolution techniques that estimate cell type composition under each spot (Li et al., 2022; Miller et al., 2022; Li et al., 2023) could be used to address this problems.

We have attempted to filter out genes that might mislead phylogenetic inference. However, among the selected highly variable genes, there still could be genes whose expression reflect their biological function or the impact of their environment and is not related to cell’s evolutionary history. Other approaches to filtering genes, such as finding subsets of genes consistent with a single phylogeny (Strimmer and Von Haeseler, 1997) or filtering saturated genes (Ducĥene et al., 2022) could improve phylogenetic reconstruction.

We marginally explored different expression evolution modelling approaches, focusing on the ordinal model for discretised data and the Brownian motion model for continuous counts. Future studies would benefit from considering other models of discrete character evolution, such as the Lewis Mk model (Lewis, 2001) that allows instantaneous evolution of any category to any other category or more general models that additionally allow unequal transition rates between categories or more complex continuous models, e.g., Ornstein–Uhlenbeck process (Hansen, 1997). It is surprising that our analyses where continuous counts were modelled to evolve under the Browning motion model (Felsenstein, 1973) produced considerably poorer results compared to the analyses with discretised data. A possible explanation could be that we did not transform the exponentially distributed counts to satisfy the normality assumption of the Brownian motion process in the presented analyses because initial attempts of log and square root transformations did not show improvements (data not shown). More thorough exploration of the effects of the normalisation transformations should be done. Furthermore, here we only used a strict clock model that assumes constant rate of change throughout the tree (Zuckerkandl, 1962; Zuckerkandl and Pauling, 1965). Using relaxed clock models Drummond et al. (2006) could improve the results. Finally, an error model could be developed to account for the low detection efficiency, similar to the error models developed for DNA evolution (Kozlov et al., 2022; Chen et al., 2022b), to directly incorporate the technology into the statistical inference.

To conclude, we showed that the phylogenetic relationships of malignant epithelium and benign normal epithelium from different locations can be recovered when high quality spots and HVGs are used in the analysis. There is however still many problems to address for a robust phylogenetic and phylogeography analysis of Visium ST expression data.

## 4 Methods

### 4.1 Patient and tissue description

We analysed Visium ST (v1, 3’ chemistry) data from Pinkney et al. (2024). The data was obtained from tumour and normal tissues of two patients with colorectal cancer. For patient 1, there are two serial tumour sections (we refer to these sections as Tumour1 and Tumour2) and two serial normal sections (Normal1 and Normal2). For patient 2, there are two serial tumour sections (Tumour1 and Tumour2), one normal (Normal) and an ovary metastasis (Metastasis). Tissue images were annotated by Pinkney et al. (2024) and five major tissue types were identified: malignant gland, malignant stroma, benign colon epithelium, benign stroma, benign lymphocytic aggregation. We also reanalysed 10x Chromium single cell expression data of breast cancer-derived xenografts from Moravec et al. (2023).

### 4.2 Counts matrices

We used data as prepared and pre-processed in Pinkney et al. (2024) using 10x Genomics SpaceRanger 1.2.2. They processed raw sequencing data into fastq format (spaceranger mkfastq pipeline) followed by mapping and counting reads to form spatial gene expression matrices for fresh frozen (FF) tissue (spaceranger count pipeline). For our analysis, the eight capture areas were then aggregated using spaceranger aggr pipeline. For patient 1 50% density sample and patient 2 50% density and test samples non-aggregated data were used.

### 4.3 Subsamples

We have analysed five subsamples from patient 1 and two samples from patient 2. One sample was within a single tumour section (Tumour2) of patient 2 (*test set*). The rest of the samples were across four sections of the same patient: *50% density* samples from patient 1 and patient 2, manually *selected* spots from patient 2, *highest* quality spots from patient 2, *similar* quality spots from patient 1 and patient 2.

#### 4.3.1 50% density

A sample across all sections of patient 1 (719 spots) and a sample across all sections of patient 2 (2166 spots) were obtained using the filtering algorithm from Moravec et al. (2023), that is described above (section 4.5).

#### 4.3.2 Test set

We manually selected ten spots from two malignant and one stroma regions (30 spots in total) from Tumour 2 section of patient 1. We visually assessed spots on the high quality H&E images and selected adjacent spots that cover only one type of tissue: either malignant epithelium or stroma (Supp. Figure 12). Similarly, to all analyses in this study, we treat each spot as a single cell.

#### 4.3.3 Manually selected

We manually selected malignant and benign epithelium spots from distant regions from each of the four sections of patient 2 (25 spots). Spots on two serial tumour sections were selected from the same regions on each section (Supp. Figure 21).

#### 4.3.4 Highest quality spots

We selected highest quality malignant and benign epithelium spots from four sections of patient 2 using the following procedure. Based on the pathologist annotation we split all spots in four groups:

- benign epithelium from normal section,
- benign epithelium from two tumour sections,
- malignant epithelium from two tumour sections, and
- malignant gland from metastasis section,

We removed outlier spots with unusually high expression (higher then 99% quantile) in each group. In total we removed three spots in the malignant epithelium from tumour sections and five spots from each of the three remaining groups. Then we separated benign and malignant tumour groups in two based on the sections, obtaining six groups in total. We selected five spots with highest total expression counts from each of the six groups (30 spots in total).

#### 4.3.5 Similar quality spots

To avoid correlation between differences in total counts and phylogenetic distance and also avoid the circular use of the annotation as in the highest quality sample, we selected spots that have similar total counts. We kept benign normal spots that were selected in the highest quality sample of patient 2 as they had the lowest total counts among the six groups. These spots total counts ranged between 296 and 405. Then we randomly selected five spots in each of the remaining five groups with total counts within the same range (30 spots in total).

We performed a similar procedure for patient 1. To identify the range, we chose 0.95 quantile (2381.51) of the total counts distribution on Normal 2 section as the upper bound and 1500 as the lower bound. In total 25 spots were selected, five from each group except of benign Tumour groups where only one spot from Tumour 1 and four spots from Tumour 2 were within the specified range.

### 4.4 Discretisation and normalisation

For phylogenetic analysis of expression data we considered two approaches. We either discretised expression counts or used raw counts (continuous). The distribution of the non-zero expression counts resembles exponential distribution. Applying an equal quantile discretisation to such data results in single count categories for low expressed cases, (e.g., all cases where only a single RNA molecular was detected form a separate category). Here we used a more suitable and intuitive discretisation approach developed in Moravec et al. (2023), where counts within concentrated regions of the most frequent counts are considered normally expressed and outside of these regions, abnormally expressed. In particular, first distributions of non-zero counts are centred and standardised for each gene and then combined to obtain a single count distribution. Then 60% and 90% high probability density (HPD) intervals of the combined distribution are used to split counts into categories. Such a procedure produces six categories. Zero counts form a separate category (*unexpressed*, coded as 0), counts within the 60% HPD interval are considered *normally expressed* (3), counts that are within 90% HPD but outside 60% HPD form *slightly underexpressed* (2) and *slightly overexpressed* (4) categories depending on the side they appear relative to the normally expressed, and counts outside of the 90% HPD interval on both sides are *underexpressed* (1) and *overexpressed* (5).

The categories represent uneven proportions of the data with 0 accounting for the majority of the counts (Table 1). For both patients, the left end of the 90% HPD was very close or coincided (patient 1) with the boundary of the counts range, which resulted in very few or no counts (patient 1) falling within category 1. Due to further spot and gene filtering, almost all samples only had five categories except of the test set analyses with all genes or normalised HVG, where all six categories were present. Some analyses with HVG had four categories due to small number of genes and spots. In all cases with missing categories, present categories were relabelled by shifting codes to the left to obtain consecutive numbers starting with zero.

**Table 1:**
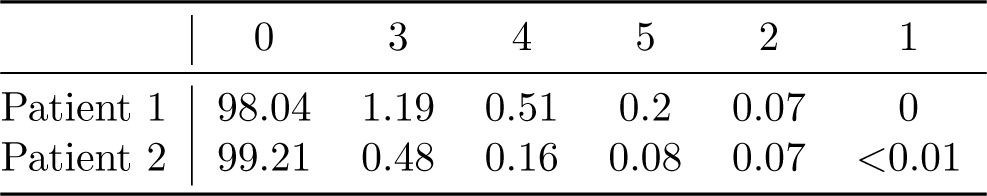
Expression category distribution in % of the expression matrix entries before gene and spot filtering.

**Table 2:** Summary of sections total count distributions.

|  | P1 Normal 1 | P1 Normal 2 | P1 Tumour 1 | P1 Tumour 2 |
| --- | --- | --- | --- | --- |
| Mean | 1005. | 1020.23 | 3157.76 | 2793.612 |
| SD | 1830.43 | 2093.66 | 4795.16 | 3636.65 |
| CV | 1.82 | 2.05 | 1.52 | 1.35 |
|  | P2 Normal | P2 Tumour 1 | P2 Tumour 2 | P2 Metastasis |
| Mean | 52.88 | 604.69 | 506.16 | 1332.77 |
| SD | 105.29 | 925.83 | 784.73 | 1707.02 |
| CV | 1.99 | 1.53 | 1.55 | 1.28 |

In the analyses of normalised and discretised data, we first normalised (divided each count in the count matrices by the total UMI of the spot) and then discretised the data. For downsampling, we used SampleUMI from Seurat v. 4.3 package (Hao et al., 2021) with the minimum total UMI count of the spots in the sample, as the target (max.umi). Downsampled values were treated as continuous.

### 4.5 Gene filtering

For 50% density samples, we used filtering algorithm proposed by Moravec et al. (2023), where spots and genes (columns and rows) in the expression matrix with smallest total expression are sequentially removed until 50% data matrix density is obtained. 2853 genes remained after filtering in patient 1 and 604 genes in patient 2.

In all other samples we either used all genes that were expressed in at least one spot of the sample (7658 genes in patient 1 similar quality sample; for patient 2: 2974 genes in similar quality sample, 3665 in manually selected sample, 6537 in highest quality sample, 4197 in test set) or only used HVGs. We identified genes as HVGs if their variance across spots in all sections of the same patient exceeded a threshold of ten (211 HVGs for patient 1 and 115 HVGs for patient 2). For the test set we considered the variance across spots in the patient 2 Tumour2 section only and lowered the threshold to two to obtain a larger subset of genes (198). The lists of HVGs for each analysis can be found in the data repository.

### 4.6 Phylogenetic analysis

In all cases with discretised data we used an ordinal model that assumes neighbouring categories can instantaneously evolve one into another with equal rates.

We performed maximum likelihood analyses using iqtree2 version 2.1.2 (Minh et al., 2020) with the ordinal model and ascertainment correction for excluding constant characters (constant zero columns). This implementation assumes equal frequencies. BEAST2 v2.6.7 (Bouckaert et al., 2019) analyses of discrete data were performed using the ordinal model with estimated frequencies and no ascertainment correction. For continuous counts, we used the Brownian motion model as implemented in contraband package v1.0.0 (Zhang et al., 2021) of BEAST2. We estimated each gene variance and root value and set covariance values to zero. In all analyses, we used a strict clock model that assumes that the overall substitution rate is equal on all branches of the tree. We assessed convergence of BEAST2 runs in Tracer v1.7.2 (Rambaut et al., 2018) with all ESS values exceeding 200. We plotted trees using ggtree R package (Yu et al., 2018; Yu, 2020).

### 4.7 Analysis of Moravec et al. (2023) data

We reanalysed single cell expression data (10x Genomics Chromium) of 58 selected cells (30% density sample) from Moravec et al. (2023) with zero counts treated as missing values. We performed two analyses, one with the original data where categories were relabelled from initial 1-5 to 0-4. In the second analysis, we only used HVGs of 58 selected cells. We calculated gene variance for unfiltered data and identified 226 HVGs (variance *>* 10).

### 4.8 Data availability

Scripts used for pre-processing of the data and estimated trees are available in https://github.com/bioDS/VSTExprPhylo repository. The Visium ST data is available upon request.

## Acknowledgments

We thank Alexei Drummond and Allen Rodrigo for there helpful discussions and suggestions throughout this research. AG was partially supported by Royal Society Te Apārangi through a Rutherford Discovery Fellowship (UOC1702) and a Marsden grant (21-UOC-057), and a Data Science Programmes grant (UOAX1932). AG and ASG were supported by Ministry of Business, Innovation, and Employment of New Zealand through an Endeavour Smart Ideas grant (UOOX1912). HP was supported by a University of Otago Doctoral Scholarship.

## 6 Supplementary material

## 6.1 Supplementary tables

## 6.2 Supplementary figures

**Figure 8:**
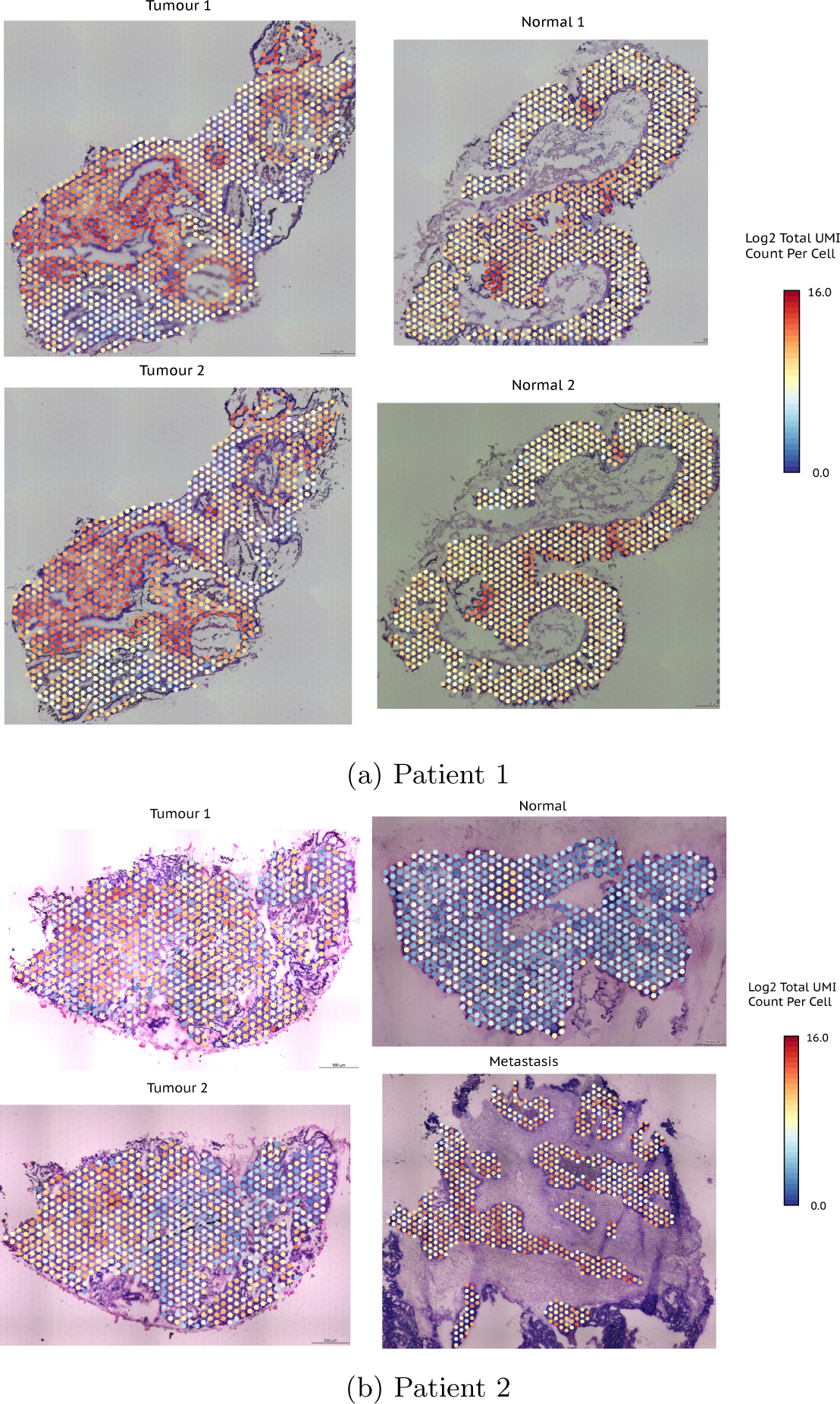
Sections images with capture spots coloured by their total UMI counts. (Top) Patient 1. Two serial tumour and two serial normal sections. (Bottom) Patient 2. Two serial tumour sections, one normal, and one metastasis.

**Figure 9:**
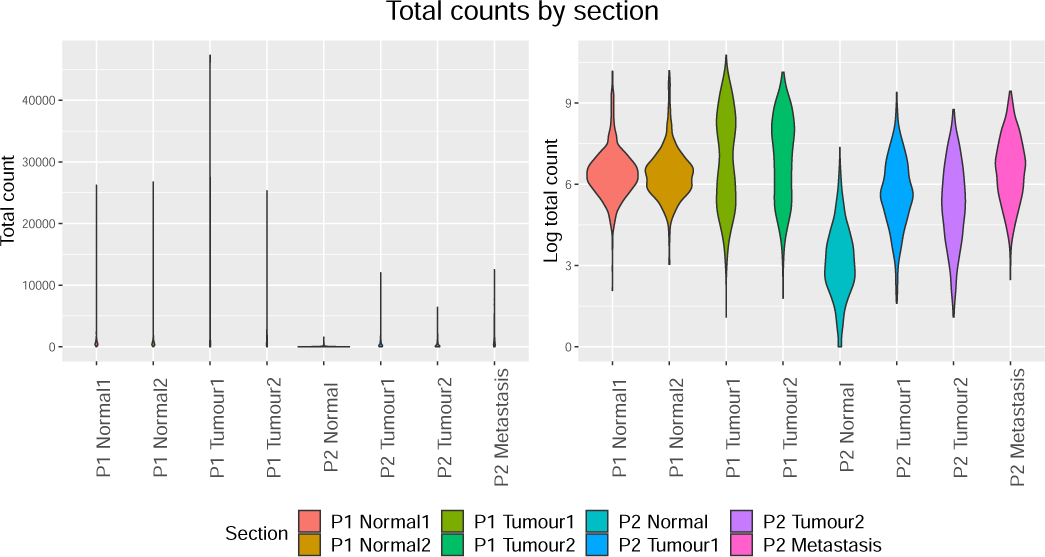
(Left) Absolute total expression (UMI) counts for different sections P1 = patient 1 and P2 = patient 2. (Right) log-transformed counts.

**Figure 10:**
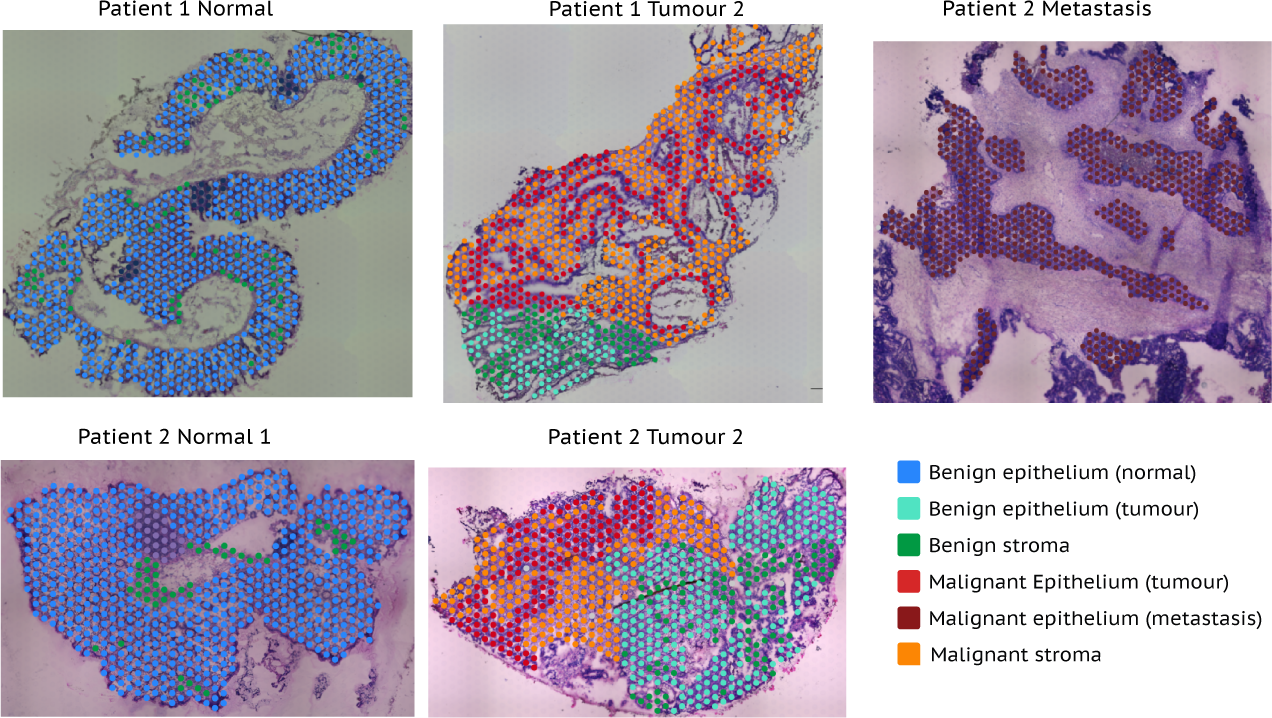
Section annotation

**Figure 11:**
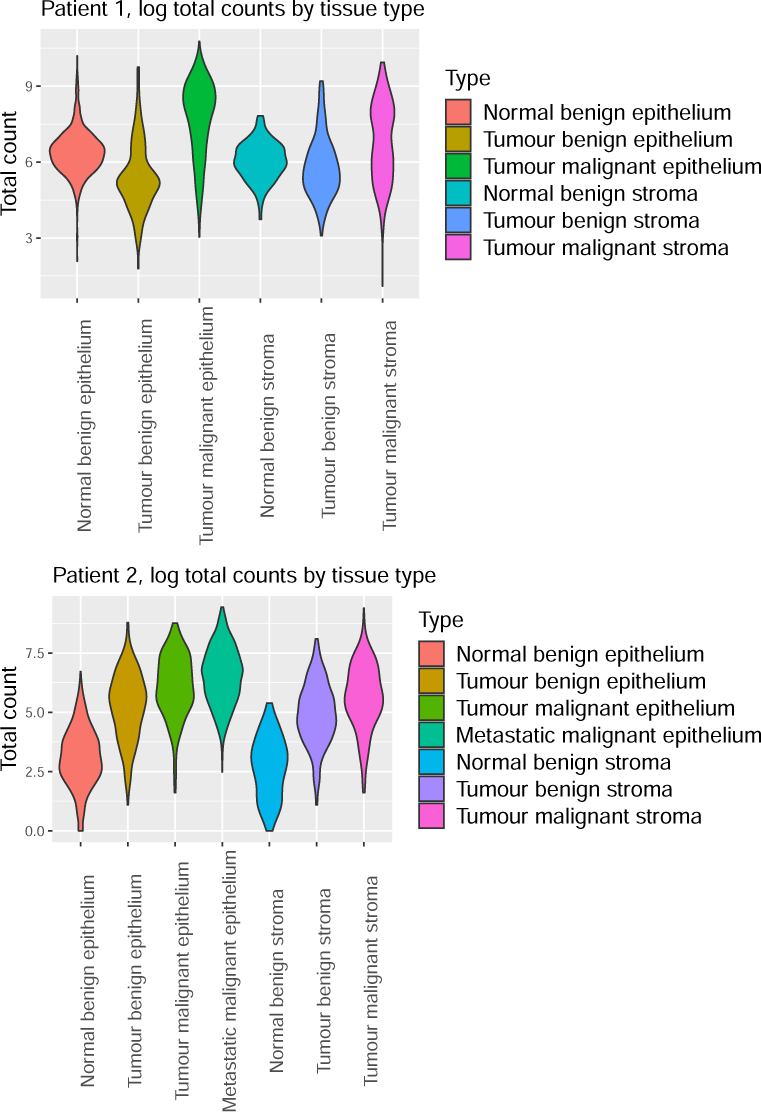
log-transformed total UMI counts for different tissue types in patient 1 (top) and patient 2 (bottom).

**Figure 12:**
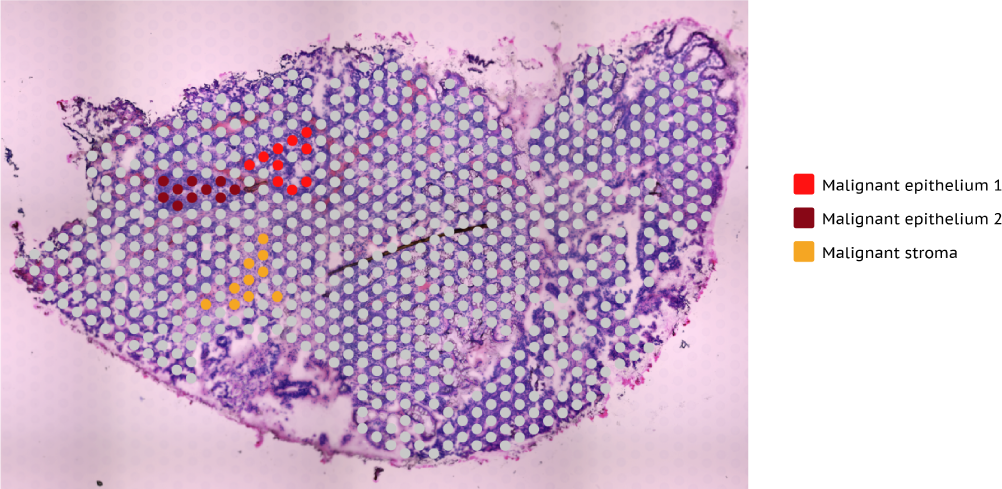
Test set from Tumour 2 patient 2. Two malignant epithelium regions and one malignant stroma region were selected.

**Figure 13:**
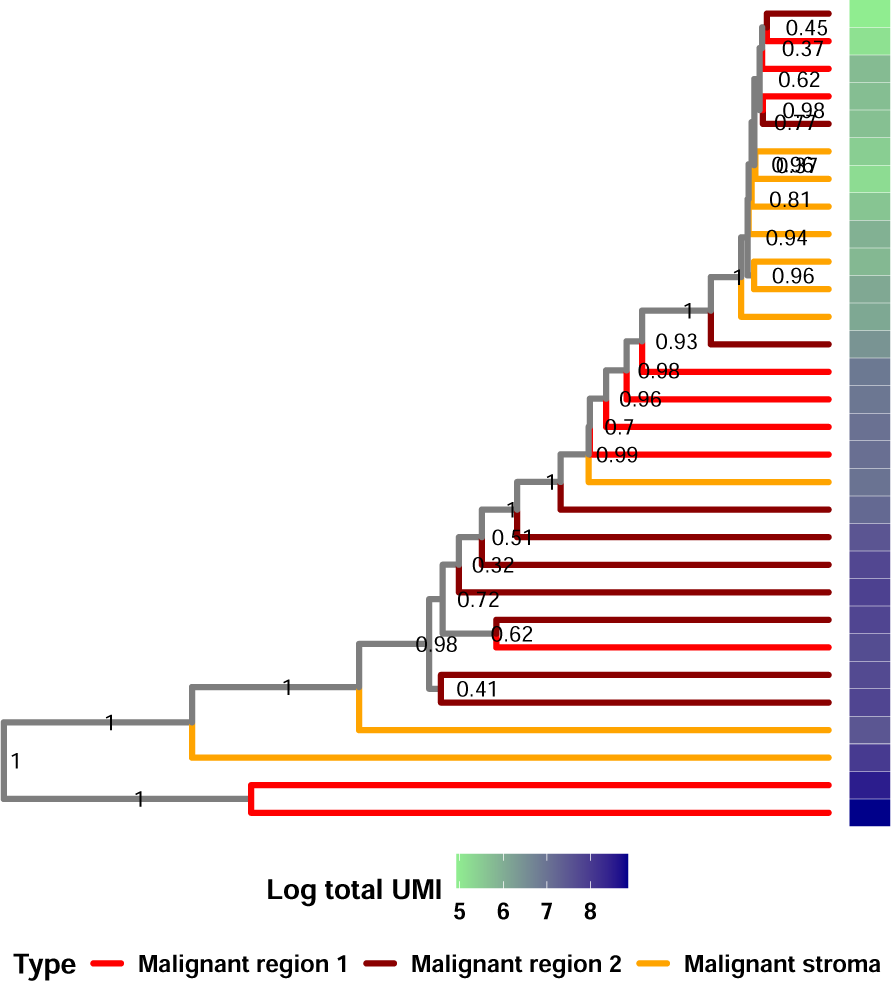
MCC tree for Bayesian analysis of discretised data of all expressed genes of the test set sample.

**Figure 14:**
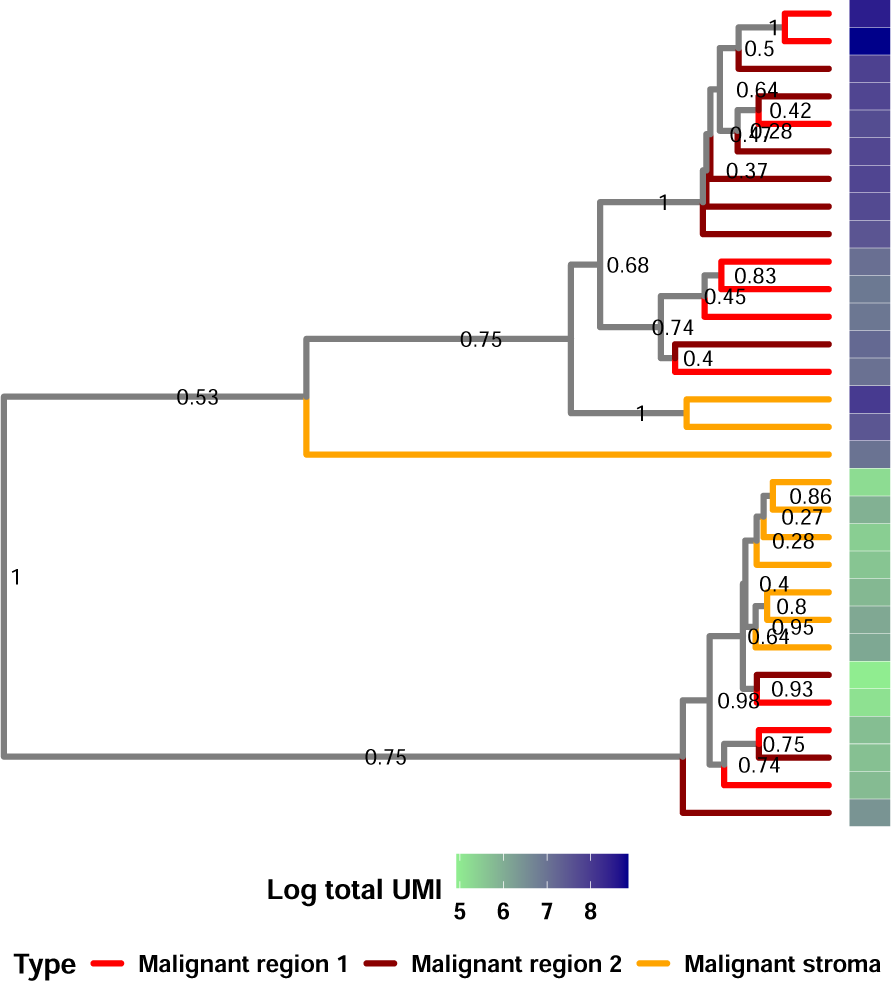
MCC tree for Bayesian analysis of discretised data of HVGs of the test set sample.

**Figure 15:**
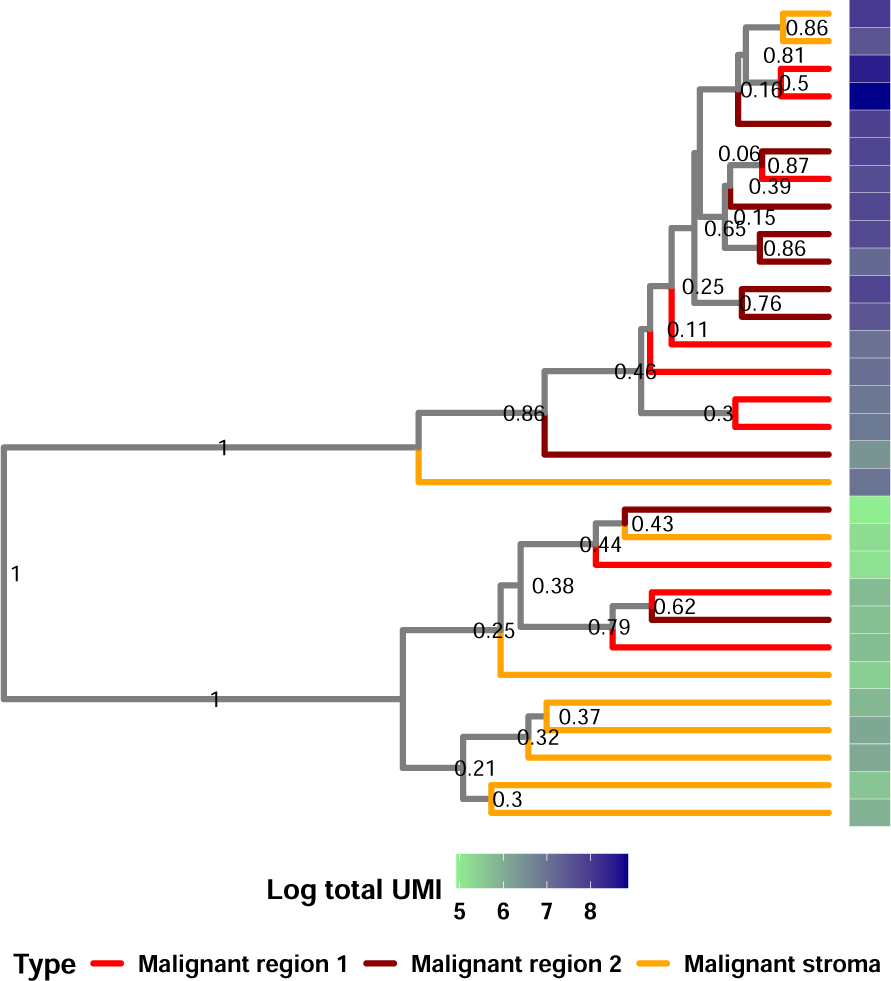
MCC tree for Bayesian analysis of normalised by spot and discretised data of HVGs of the test set sample.

**Figure 16:**
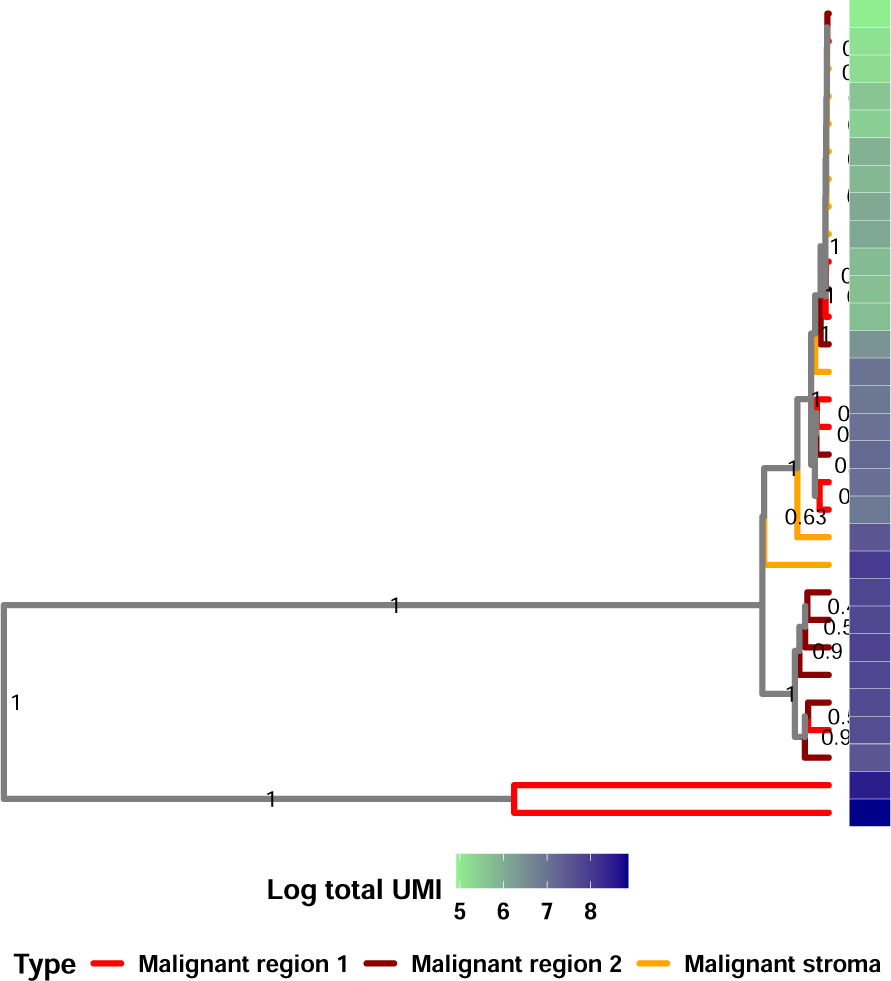
MCC tree for Bayesian analysis of continuous counts of HVGs of the test set sample.

**Figure 17:**
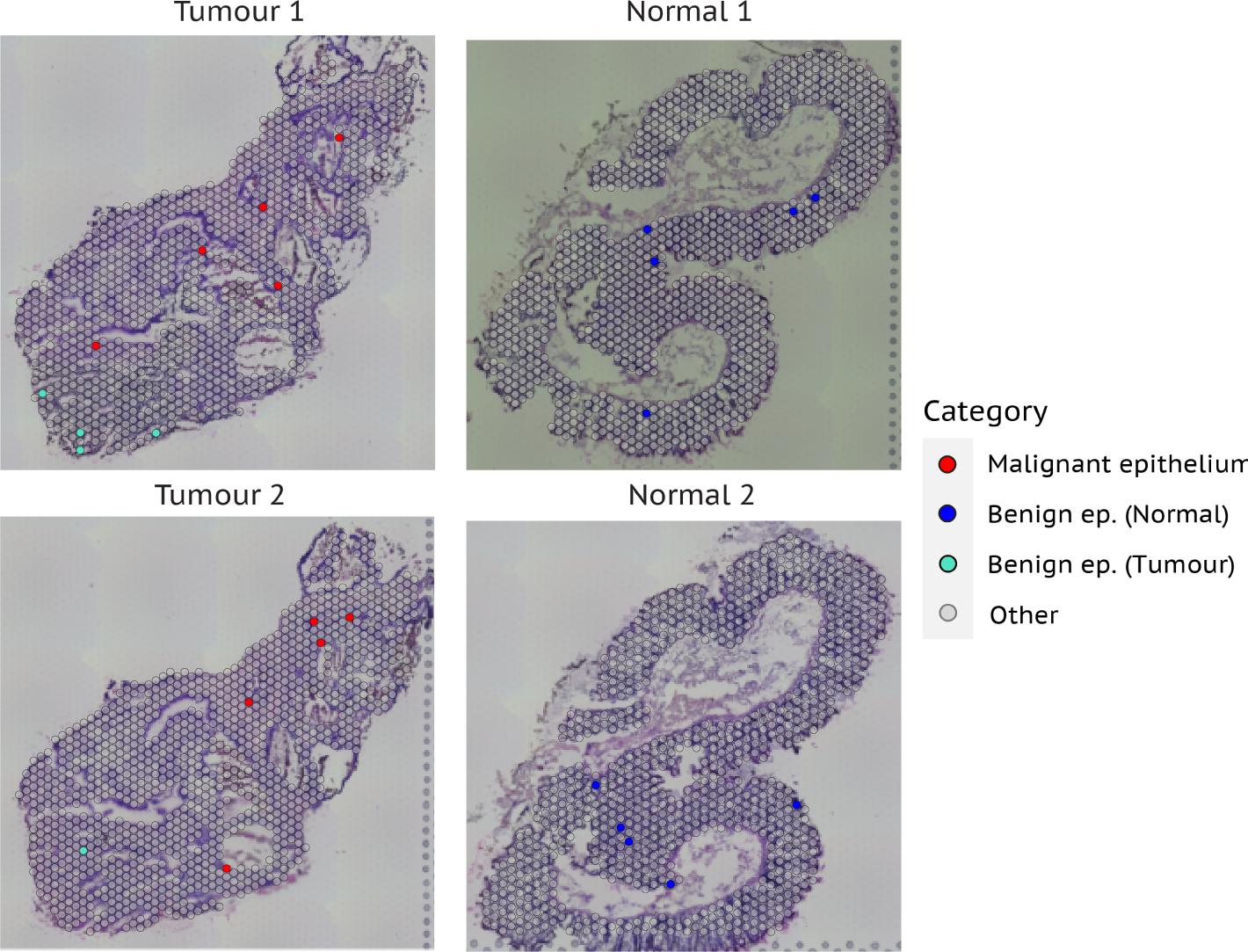
Similar quality spots, patient 1.

**Figure 18:**
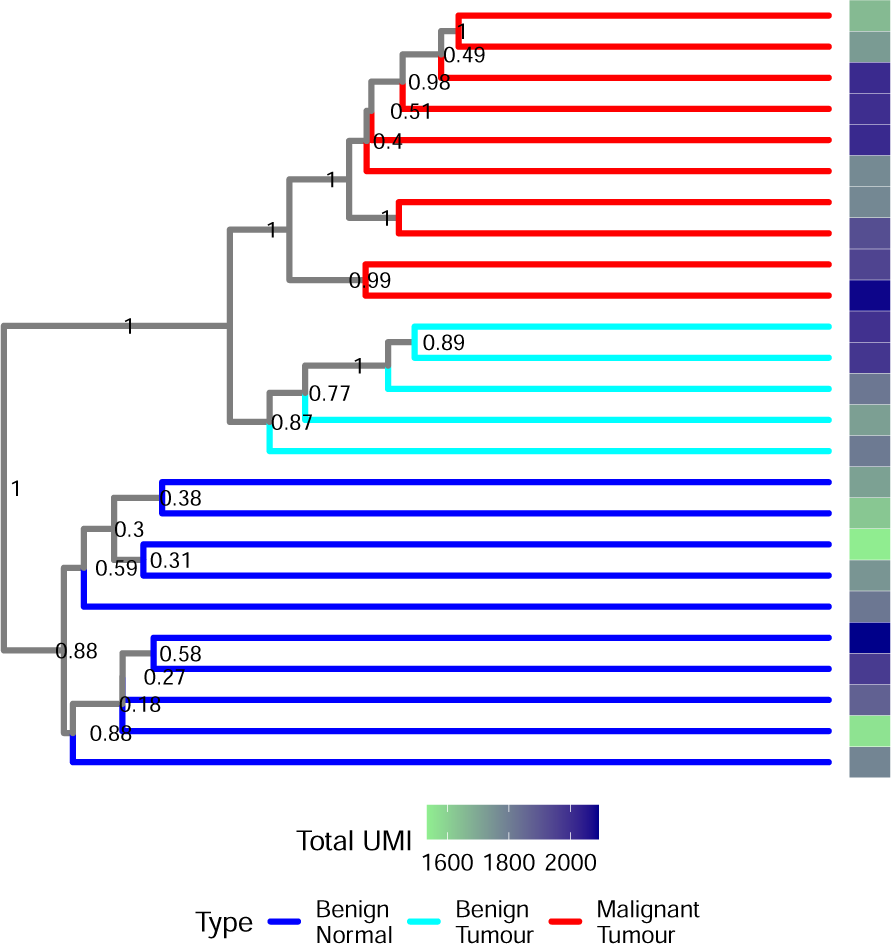
MCC tree for Bayesian analysis of discretised counts of all expressed genes for the similar quality sample from patient 1.

**Figure 19:**
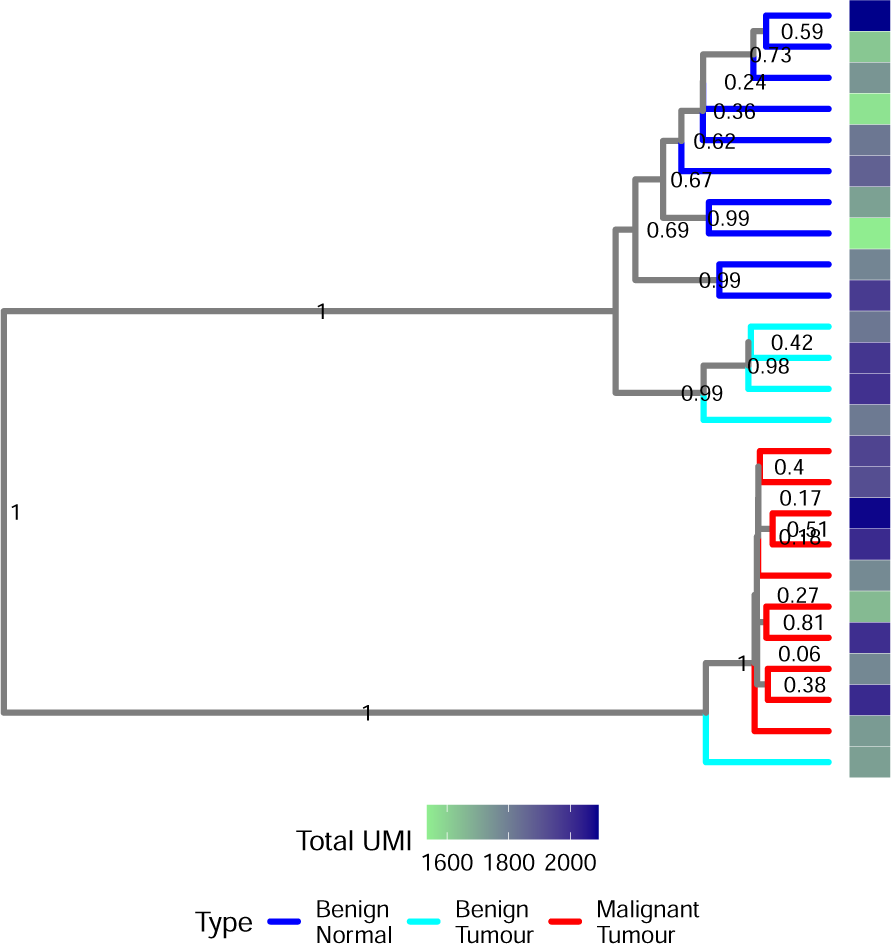
MCC tree for Bayesian analysis of normalised and discretised counts of HVGs for the similar quality sample from patient 1.

**Figure 20:**
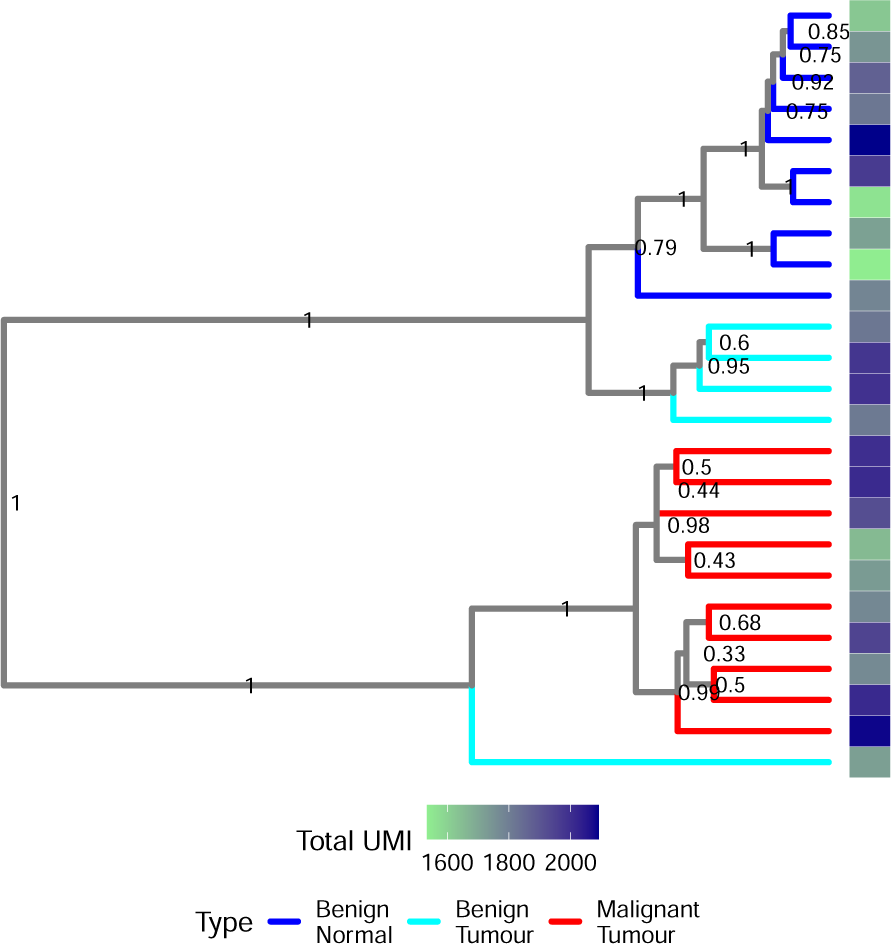
MCC tree for Bayesian analysis of continuous counts of HVGs for the similar quality sample from patient 1.

**Figure 21:**
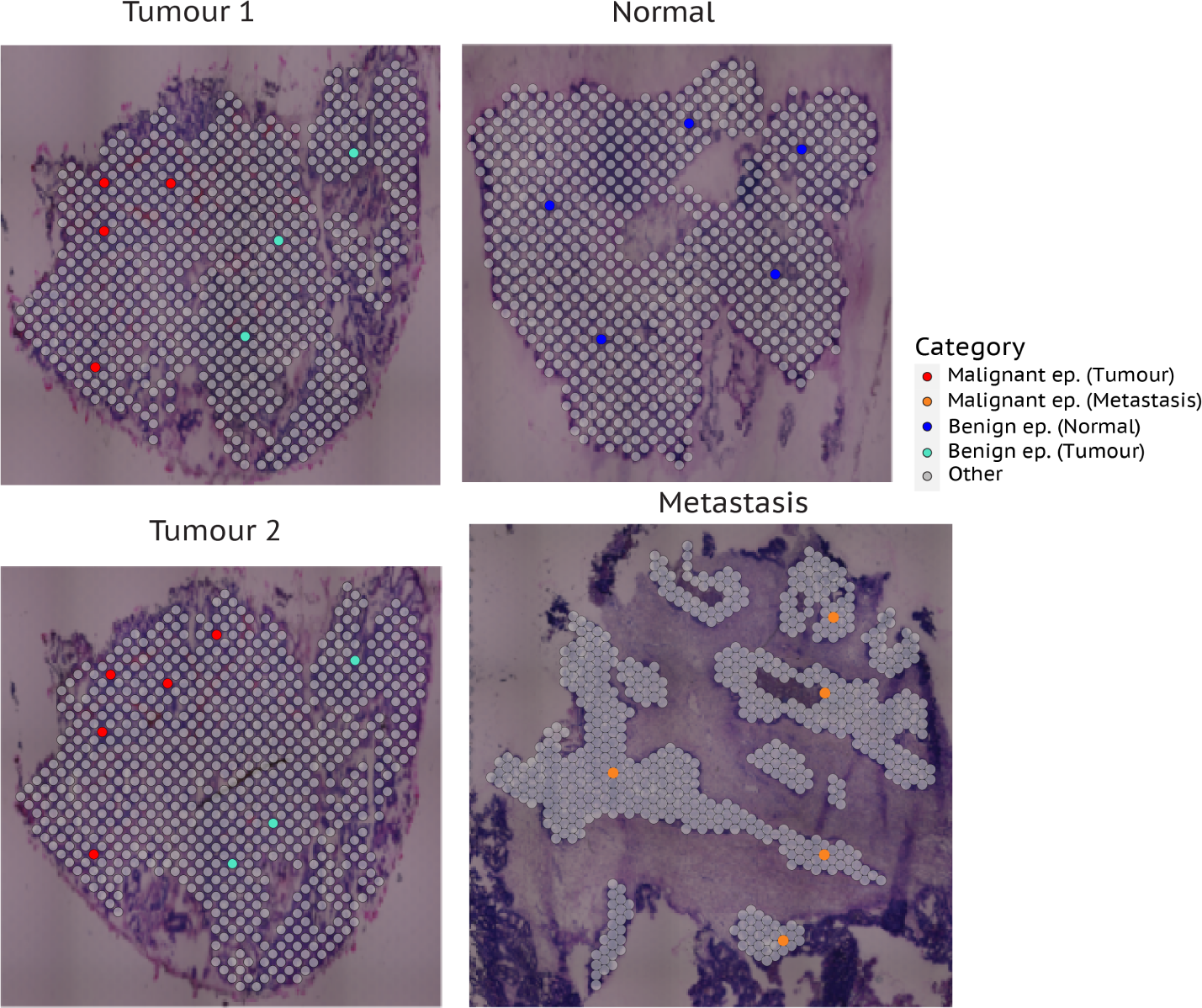
Manually selected spots, patient 2

**Figure 22:**
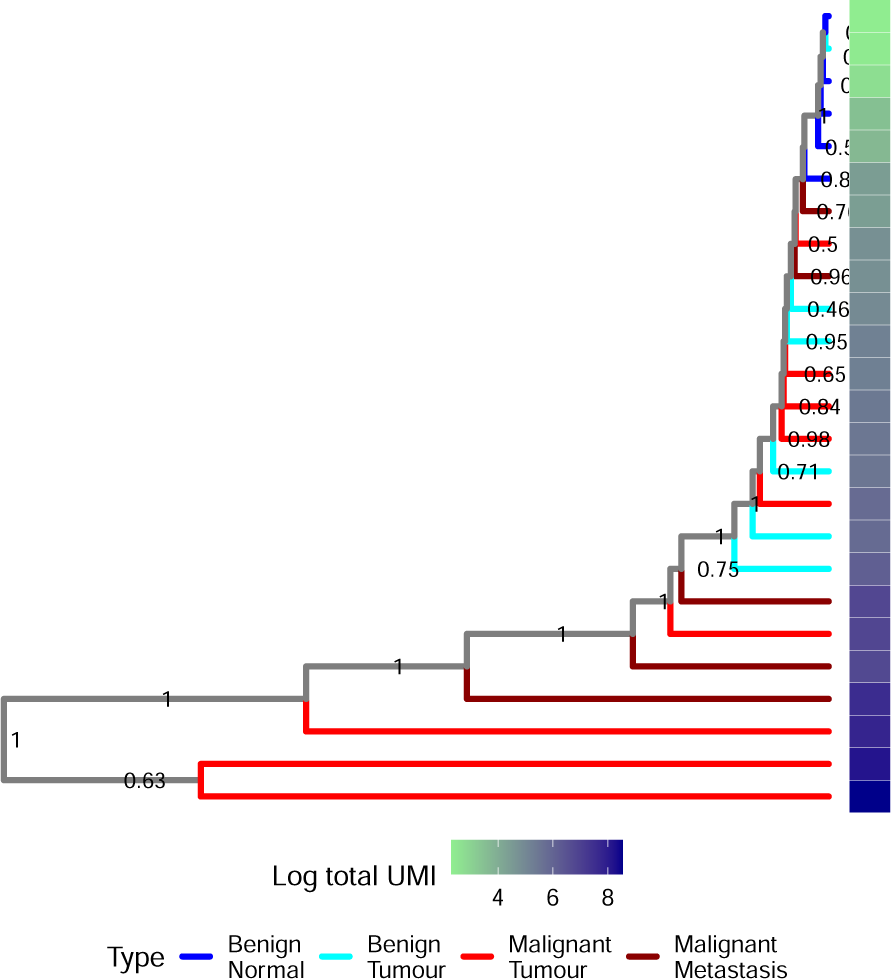
MCC tree for Bayesian analysis of discretised counts of all expressed genes for the manually selected sample from patient 2.

**Figure 23:**
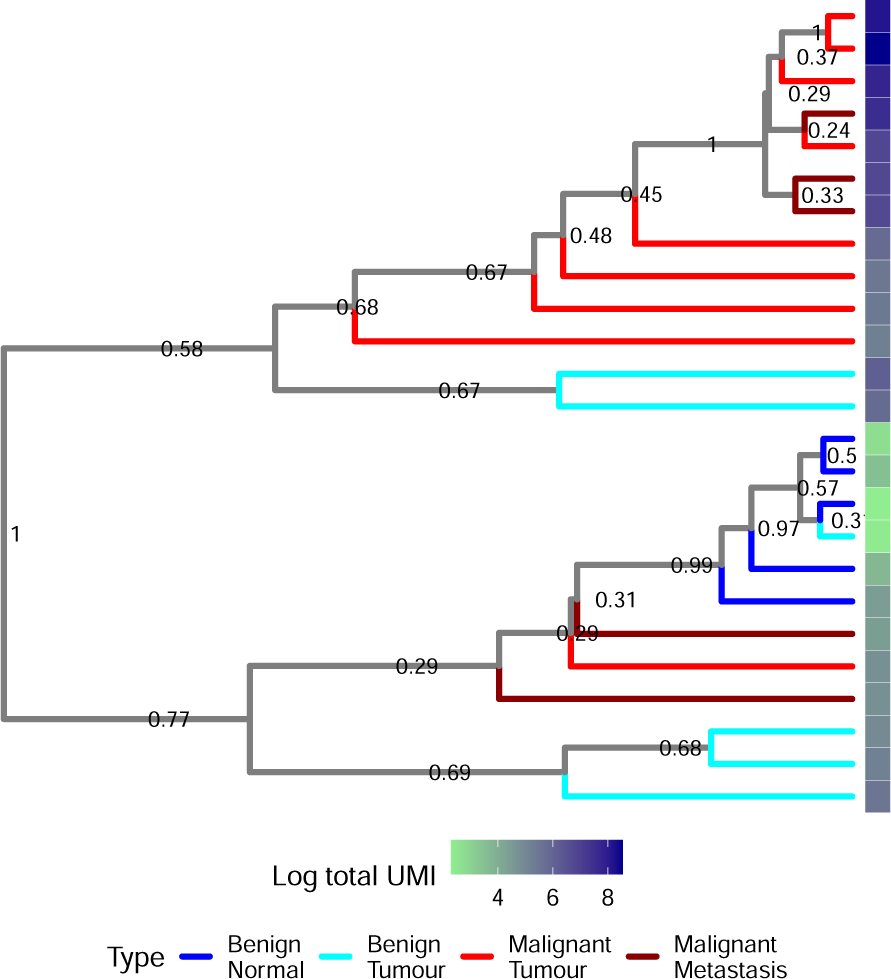
MCC tree for Bayesian analysis of normalised and discretised counts of HVGs for the manually selected sample from patient 2.

**Figure 24:**
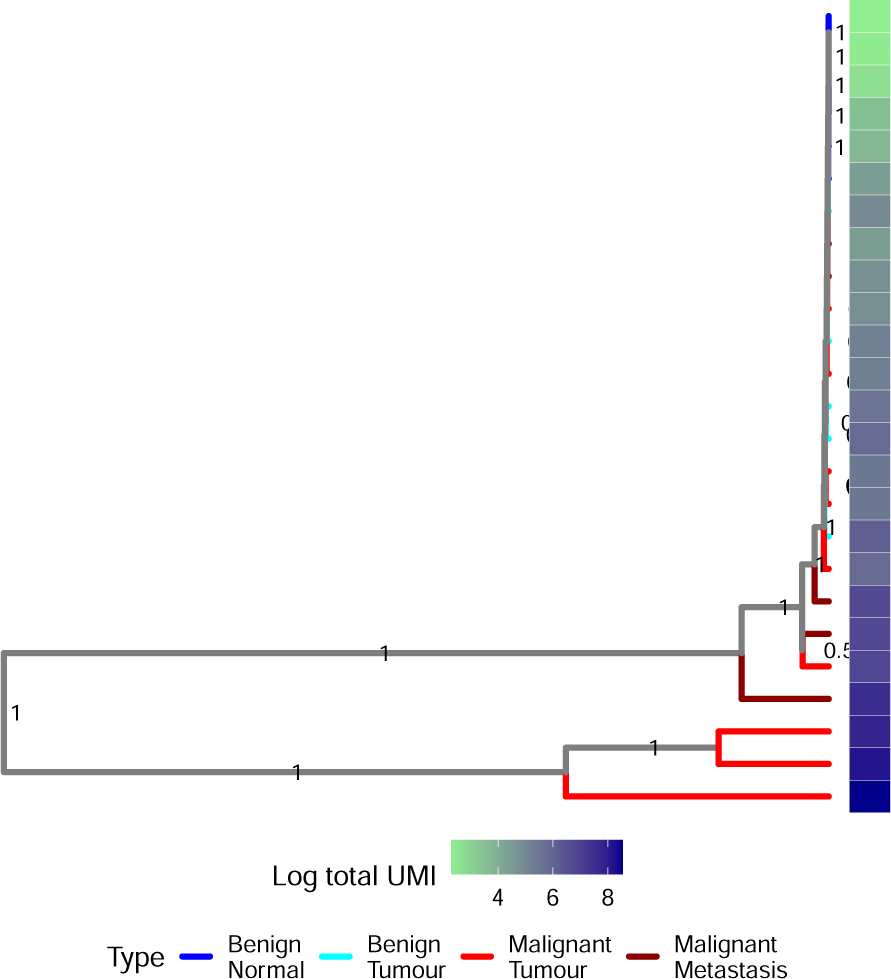
MCC tree for Bayesian analysis of continuous counts of HVGs for the manually selected sample from patient 2.

**Figure 25:**
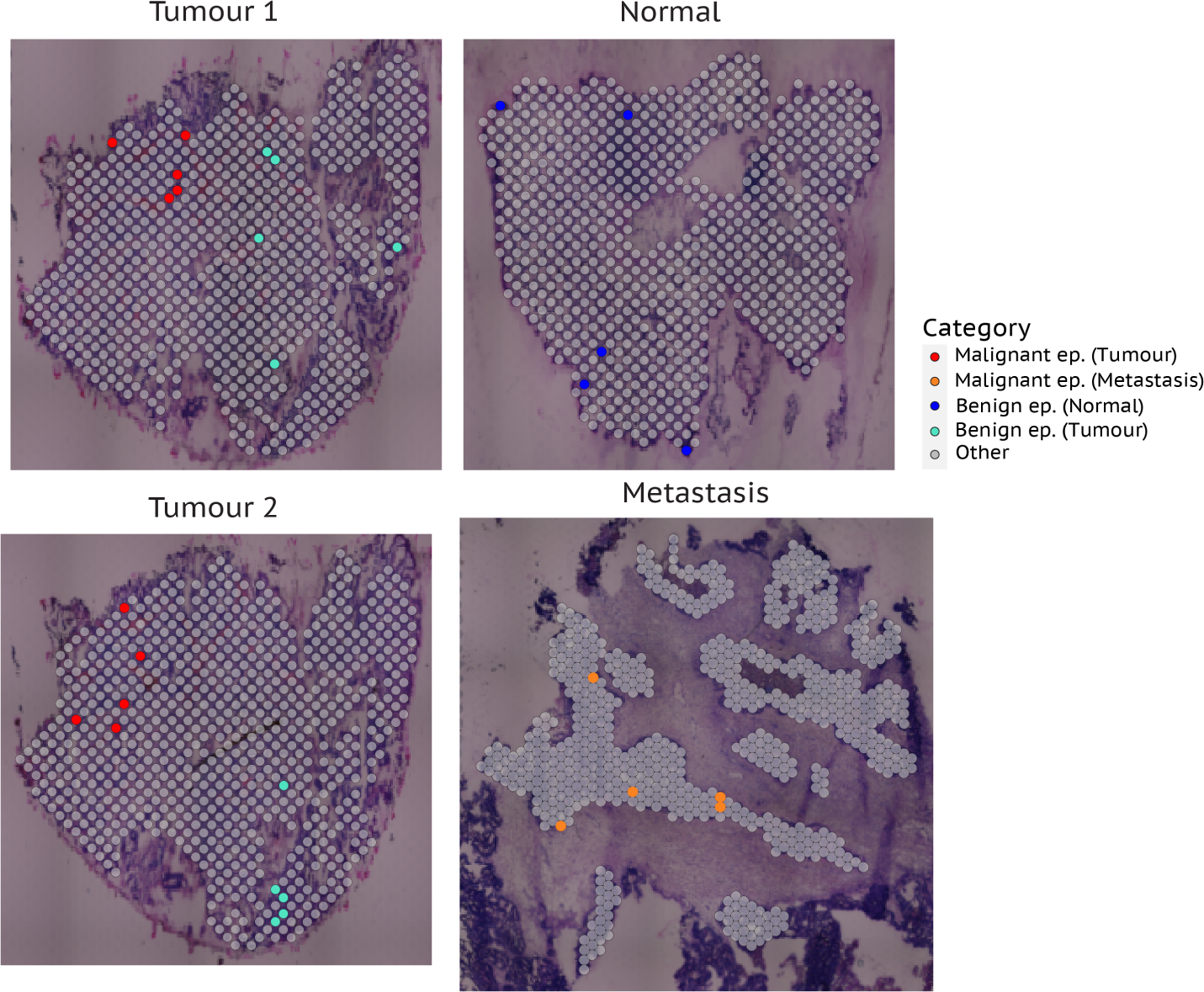
Highest quality spots, patient 2

**Figure 26:**
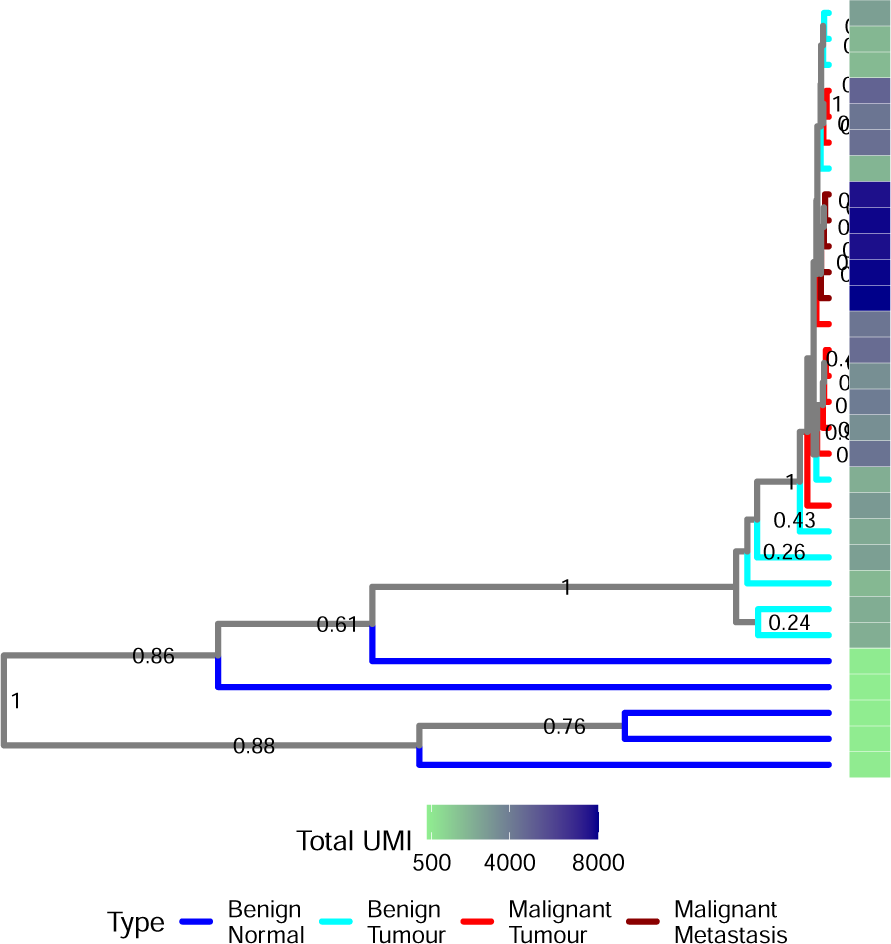
MCC tree for Bayesian analysis of normalised and discretised counts of HVGs for the high quality sample from patient 2.

**Figure 27:**
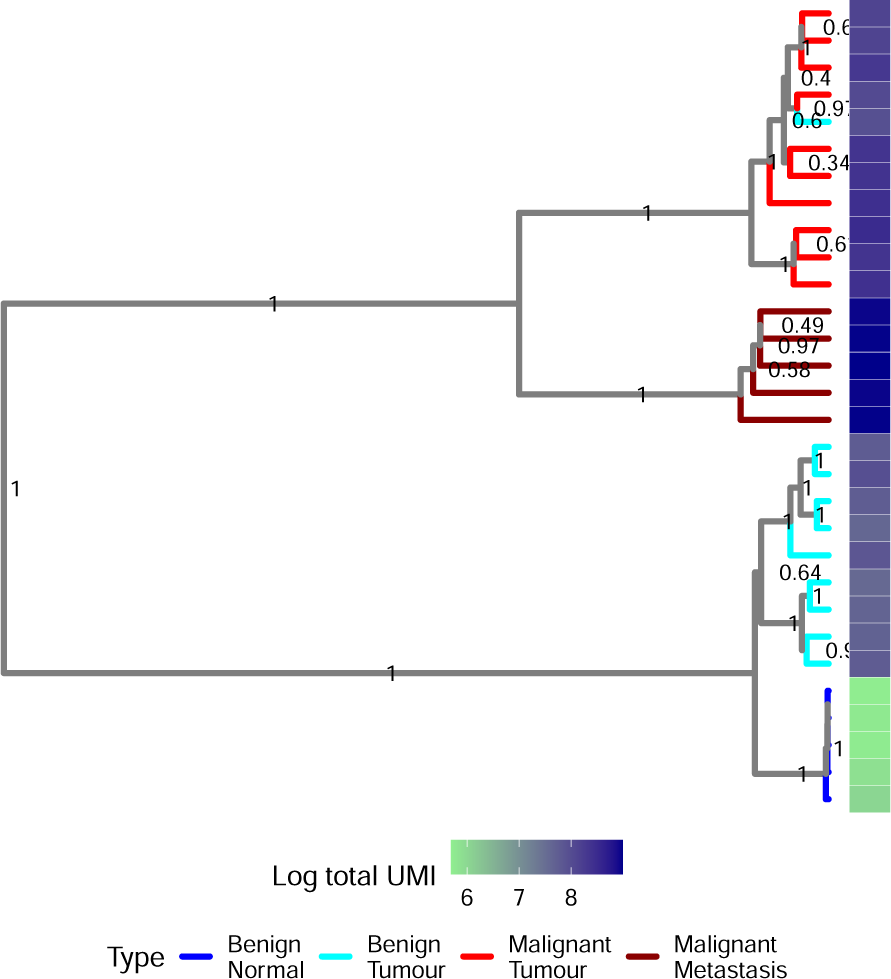
MCC tree for Bayesian analysis of continuous counts of HVGs for the high quality sample from patient 2.

**Figure 28:**
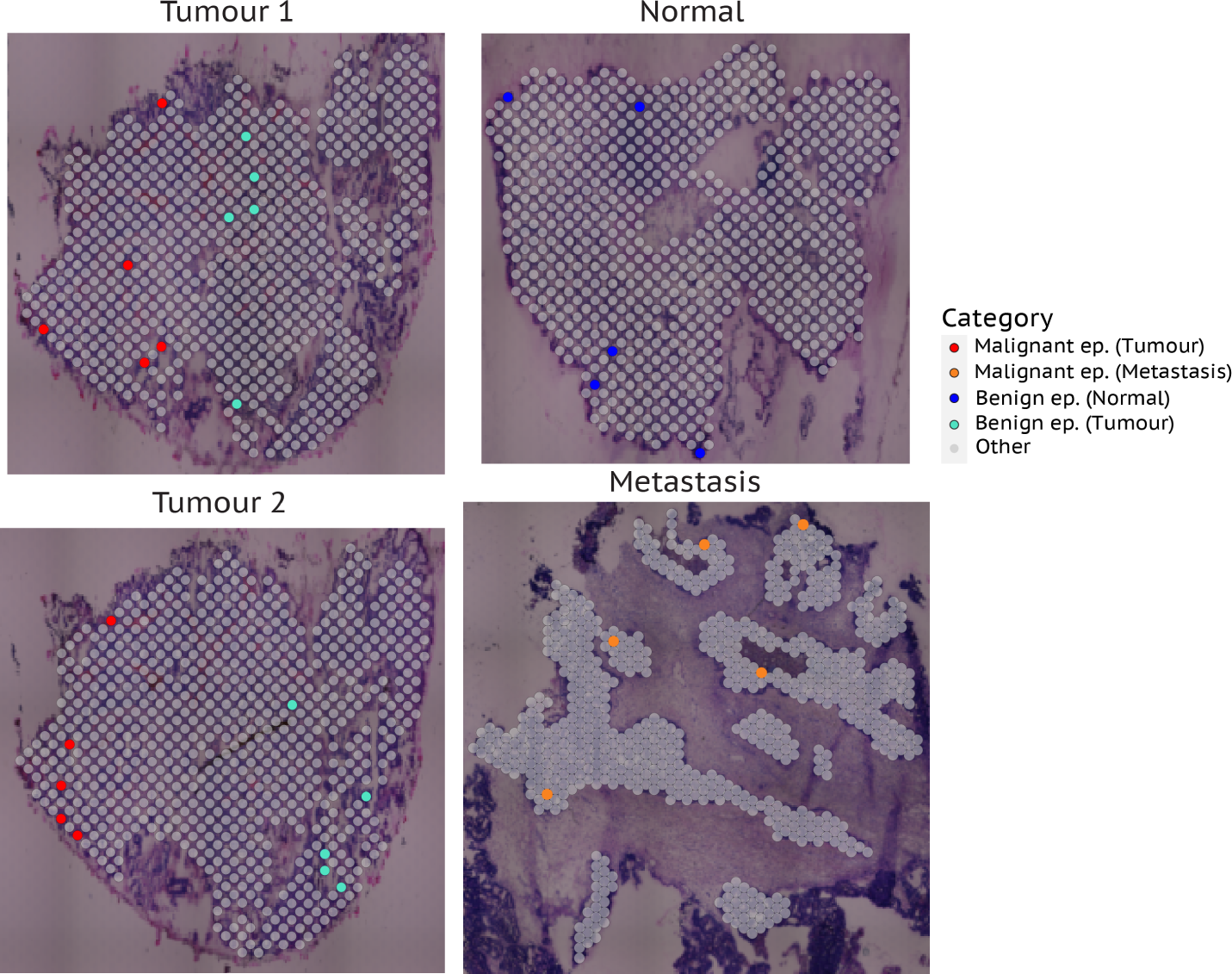
Similar quality spots, patient 1

**Figure 29:**
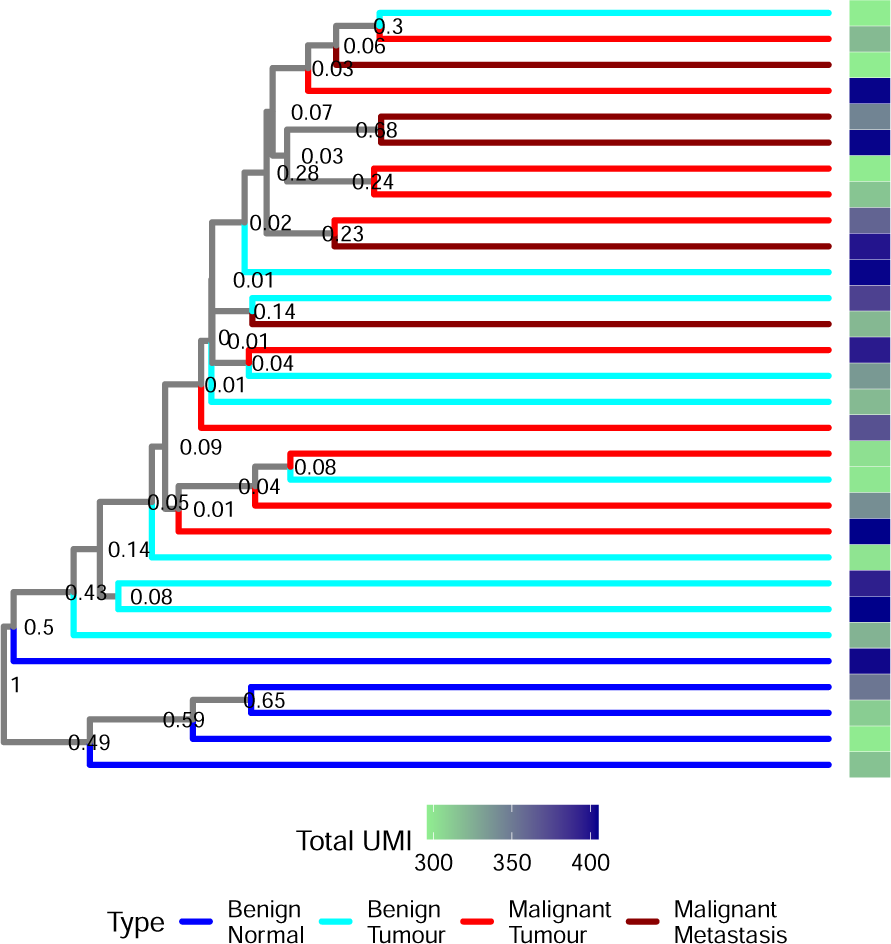
MCC tree for Bayesian analysis of discretised counts of all expressed genes for the similar quality sample from patient 2.

**Figure 30:**
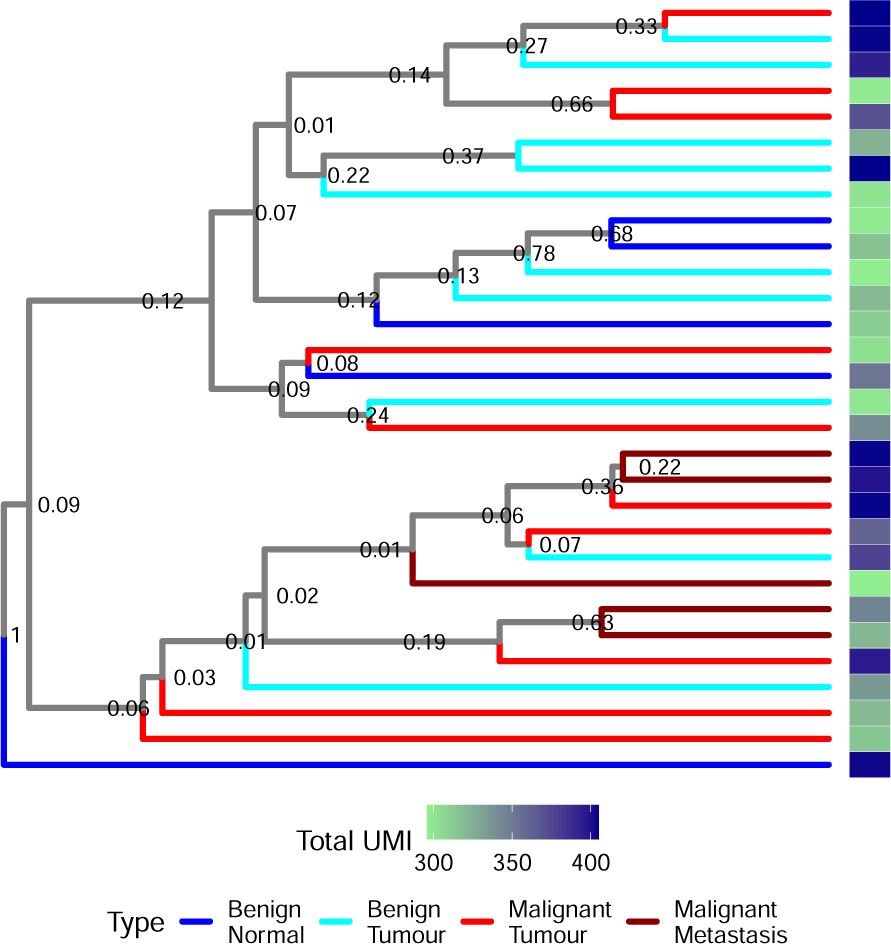
MCC tree for Bayesian analysis of normalised and discretised counts of HVGs for the similar quality sample from patient 2.

**Figure 31:**
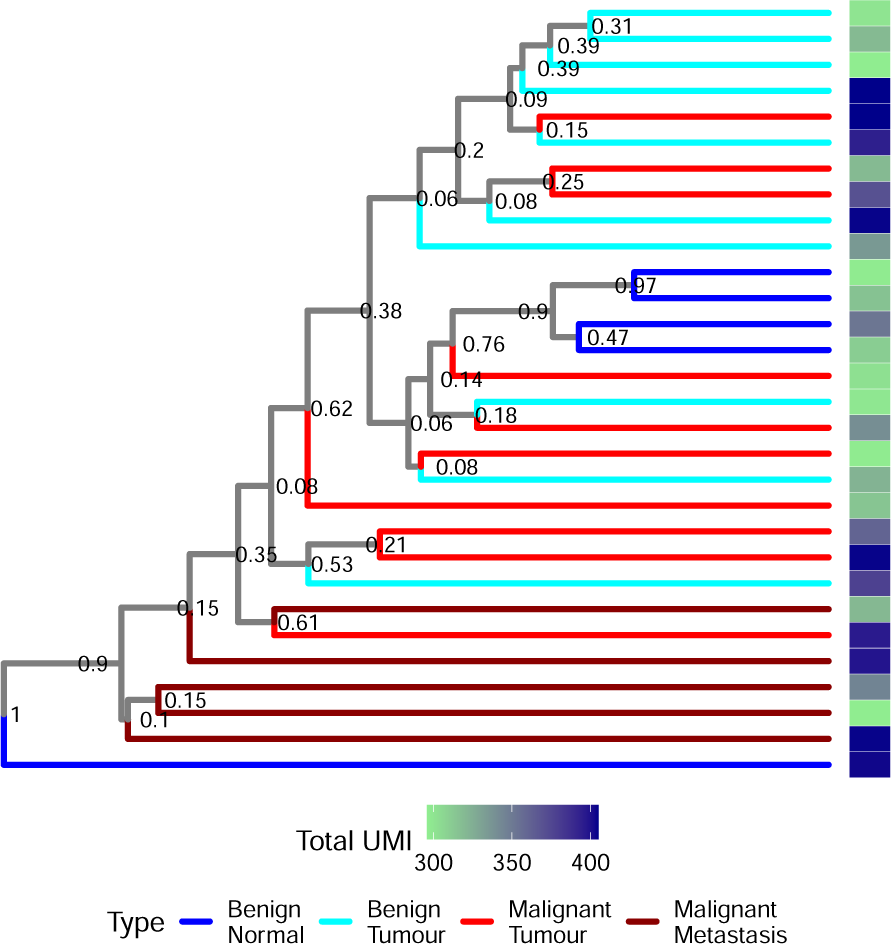
MCC tree for Bayesian analysis of continuous counts of HVGs for the similar quality sample from patient 2.

**Figure 32:**
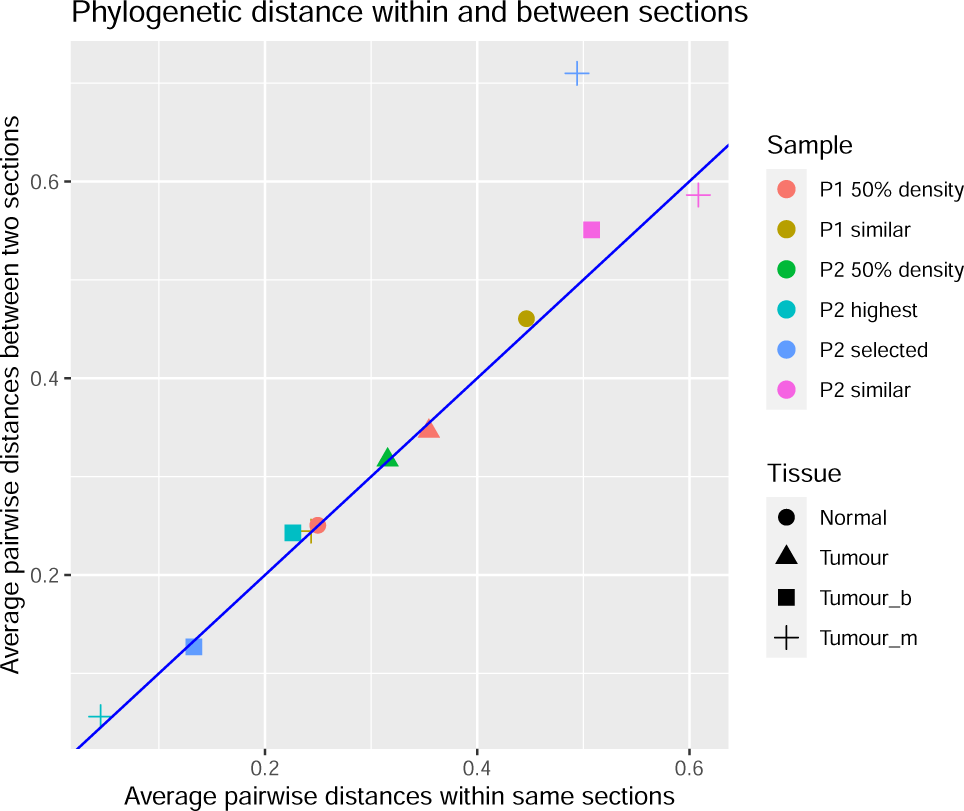
Average phylogenetic distances (normalised by the tree height or longest path for unrooted trees) within and between serial sections on maximum likelihood trees of 50% samples and MCC trees of discretised HVG analyses of four across sections samples. For all combinations of pairs of serial sections and tissue/section types, we calculated the average of phylogenetic distances between all pairs of spots from different serial sections (Y-axis). For all combination of serial sections and tissue/section types, we calculated the average of phylogenetic distances between all pairs of spots within the same section and then took the average for the two serial sections (X-axis). The blue line is a diagonal. Patient 1 similar sample benign tissue from Tumour sections case is not shown as there was only single benign spot on Tumour 1 section in this sample.

